# Novel Viroid-like RNAs Naturally Infect a Filamentous Fungus

**DOI:** 10.1101/2022.08.03.502636

**Authors:** Kaili Dong, Chuan Xu, Ruiying Lv, Ioly Kotta-Loizou, Jingjing Jiang, Linghong Kong, Shifang Li, Ni Hong, Guoping Wang, Robert H. A. Coutts, Wenxing Xu

## Abstract

Viroids have been found to naturally infect only plants, resulting in big losses for some crops, but whether viroids or viroid-like RNAs naturally infect non-plant hosts remains unknown. Here we report the existence of a set of exogenous, single-stranded circular RNAs, ranging in size between 157-450 nucleotides (nt), isolated from the fungus *Botryosphaeria dothidea* and nominated Botryosphaeria dothidea circular RNAs (BdcRNAs). BdcRNA(s) replicate autonomously in the nucleus *via* a rolling-circle replication mechanism following symmetric pathways with distribution patterns depending on strand polarity and species. BdcRNAs can modulate to different degrees specific biological traits (e.g., alter morphology, decrease growth rate, attenuate virulence, and increase or decrease tolerance to osmotic stress and oxidative stress) of the host fungus by regulating related metabolic pathways. Overall, BdcRNA(s) have genome characteristics similar to those of viroids and exhibit pathogenic effects on the fungal hosts. These novel viroid-like RNAs infecting fungi are proposed to be termed as mycoviroids. BdcRNA(s) may be regarded as additional inhabitants at the frontier of life in terms of genomic complexity, and represent a new class of acellular entities endowed with regulatory functions, and novel epigenomic carriers of biological information.

**Significance statement:** Several viroids have been transfected into unicellular and filamentous fungi to assess whether they can replicate, but no natural infections of fungi with viroid or viroid-like RNAs have been reported before. Here we describe a set of exogenous circular RNAs (cRNAs) in a phytopathogenic fungus. These cRNAs display molecular and biological features which might represent a new class of viroid-like cRNAs endowed with regulatory functions, and novel epigenomic carriers of biological information. This is the first report of infectious viroid-like RNAs (or exogenous small cRNAs) in a life kingdom (fungi) other than plants. We also present a subcellular analysis of cRNAs in a fungus for the first time and provide useful understanding in how cRNAs replicate, move, and are distributed in fungal cells.

## 1. Introduction

Viroids are small pathogenic single-stranded (ss) protein-coding RNAs varying in size from 246 to 401 nt, which replicate autonomously following inoculation of higher plants ^[1]^. Viroids are classified into two families, *Pospiviroidae* and *Avsunviroidae*, based on secondary structure, ribozyme activity, subcellular location, genome organization, replication, and phylogenetic relationships ^[1]^. Members of the *Pospiviroidae* family adopt a rod-like secondary structure with five structural/functional domains, including a central conserved region (CCR), and are localized and replicate in the nucleus ^[1b]^. Members of the *Avsunviroidae* family adopt a branched or quasirod-like secondary structure, lack a CCR, form hammerhead ribozymes (HHRz), and are localized and replicate in chloroplasts ^[1]^. Viroids replicate *via* a rolling circle mechanism in three steps, RNA transcription, processing and ligation; they follow either an asymmetric pathway catalyzed by host enzymes (RNA polymerase, RNase, and RNA ligase) for members of the *Pospiviroidae*, or a symmetric pathway for members of the *Avsunviroidae* that are self-cleaved by HHRz, which functionally substitute for RNase during replication ^[1a]^. With a minimal genome size and simple structure, viroids are giants in terms of functional versatility in the RNA world, can triumph over cellular RNA degradation machineries and further enlist host-encoded factors for self-replication in specific subcellular compartments and trafficking through plants ^[2]^.

Viroids infect horticultural plants, including vegetables, fruits, and ornamentals, and cause devastating diseases in some crops ^[1b]^. To date, viroids (or viroid-like RNAs) have been found to naturally infect only plants, although several studies demonstrated that they can also replicate in yeast (unicellular fungi), filamentous fungi, oomycetes, and cyanobacteria following artificial inoculation ^[3]^. Whether viroids or viroid-like RNAs naturally infect other organisms apart from plants is not known. If they do, their molecular and biological traits, phylogenetic relationships with the viroids that infect plants, and interactions with their host is of great interest.

Phytopathogenic fungi cause numerous diseases in plants resulting in significant, often catastrophic, annual economic crop losses, and have led to human famine throughout history. Some fungi play crucial roles in ecosystems and nutrient recycling as decomposers of dead plant material and are used in the food industry and for antibiotic production ^[4]^. Other fungi cause animal and human diseases often resulting in death and disability. ^[5]^. Fungi connect diverse life kingdoms and accommodate diverse microbe types: fungi may be parasites of plants, animals (including humans), and other fungi, while also harboring viruses (known as mycoviruses or fungal viruses), protists, bacteria, and prions. Fungi together with these microbes play important roles in influencing other life kingdoms, and therefore understanding, controlling, and applying fungal communities with the aid of symbiotic organisms attracts major theoretical and practical interest.

*Botryosphaeria dothidea* (Moug.: Fr.) Cesati & De Notaris (anamorph *Fusicoccum aesculi* Corda) is an important phytopathogenic fungus with a worldwide distribution that infects numerous tree species including apple, pear, and grape causing symptoms including die-back, stem and shoot blight, gummosis, canker and fruit rot ^[6]^. It is known that most fungal species (including *B. dothidea*) are highly divergent in their biological traits including morphology, growth rate, and virulence in the same host; even if their genetic background is identical, as is the case for conidium- or protoplast generated sub-isolates of the same parental strain. These changes, which are beyond our understanding based on current genetic knowledge, might be elicited by certain as yet unknown regulatory factors or biological agents.

Here, we have characterized a set of exogenous circular RNAs (ecRNAs) isolated from an attenuated phytopathogenic strain of *B. dothidea* and found that they display molecular and biological features similar to those of viroids, and modulate to different degrees specific biological traits (e.g., growth rate, morphology, and virulence) of the host fungus by regulating related metabolism pathways. These ecRNAs appear to cause alterations to the biology of fungi and possibly other cellular organisms.

## 2. Results

### 2.1 A phytopathogenic fungus harbors a complex pattern of dsRNAs and circular RNAs

Three *B. dothidea* strains, XA-3, MAO-1, and MAO-2, with differing morphology and virulence, were isolated from apple branches (*Malus domestica* Borkh. cv. ‘Fuji’). Strain XA-3 was avirulent or weakly virulent on apple or pear fruits eliciting lesions < 1.4 cm in diameter as compared to those caused by the MAO-1 and MAO-2 strains which were > 3.0 cm in diameter (Figure 1A, panel II and IV). Strain XA-3 had a similar morphology and growth rate to strain MAO-1 on potato dextrose agar (PDA; Figure 1A, panel I and III). Following culture on PDA XA-3 and MAO-1 colonies were initially white but subsequently turned grey, with a flat surface and dense and cotton-like aerial mycelium. In contrast, MAO-2 had increased growth rate (of 4.0 mm per day) and formed white colonies with thin mycelia collapsed at the centre (Figure 1A, panel I and III).

**Figure 1.**
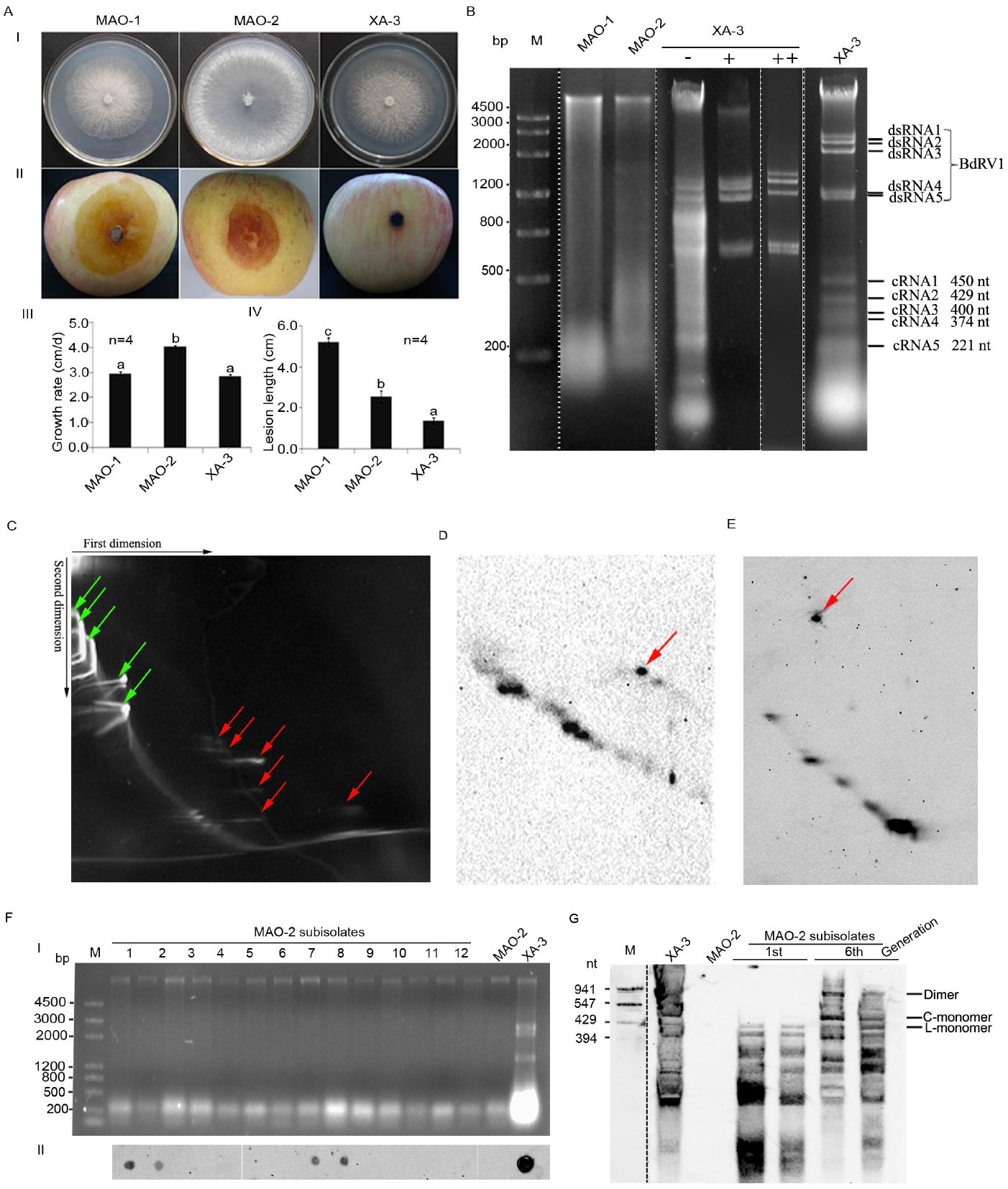
Morphology and virulence of *Botryosphaeria dothidea* strains and characterization of their associated nucleic acids. (A) Colony morphology (I), virulence on apple (M. *domestica* cv. ‘Fuji’) (II), histograms of growth rates (n=3; III) and lesion lengths on apple (n=8; IV) of MAO-1, MAO-2, and XA-3, respectively. Bars in each histogram labeled with the same letters are not significantly different (P >0.05) according to the least significant difference test (1-way ANOVA); and error bars represent standard deviation (SD). (B) Electrophoretic analysis on a 1.2% agarose gel of nucleic acid preparations from XA-3 before enzymatic treatment (-), after digestion with S1 nuclease alone (+), and after digestion with both S1 nuclease and DNase I (++); and from MAO-1 and MAO-2 without enzyme treatment. The dsRNAs 1-5 represent the genome of Botryosphaeria dothidea RNA virus 1 (BdRV1). (C) Two-dimensional polyacrylamide gel electrophoresis (2D-PAGE) analysis of nucleic acid preparations from XA-3 as showed in panel B. The green arrows indicate BdRV1 dsRNAs 1-5, followed by consecutive and traced host RNAs, and the red arrows indicate BdcRNAs, as judged by the corresponding electrophoretic positions and numbers. (D and E) Northern blot hybridization analysis of nucleic acid preparation from XA-3 after 2D-PAGE analysis of BdcRNAs 1 and 2, respectively. Both 2D-PAGE analyses are independent on this as shown in panel C. The signals refer to the circular (indicated by red arrows) and linear forms of BdcRNA1 or 2. (F) Nucleic acids extracted from protoplast-generated sub-isolates together with strains MAO-2 and XA-3 (I), and corresponding dot-blot analysis of BdcRNA2 (II). (G) Northern blot analysis of BdcRNA2 in the first and sixth generations of two positive sub-isolates chosen randomly, using transcribed BdcRNA2 fragments as the marker. C- and L-monomer refer to the circular and linear forms, respectively.

To investigate whether the presence of biological agents such as viruses were responsible for the divergent morphology and/or virulence of XA-3, nucleic acid preparations enriched in dsRNA were obtained from XA-3 mycelia and analyzed by agarose gel electrophoresis. A complex pattern of dsRNA bands was detected, including a population of elements, ranging from 1 to 2.4 Kbp in size, and smaller elements, ranging from 100 to 500 bp in size (Figure 1B). These elements were detected exclusively in the XA-3 isolate but not in the MAO-1 or MAO-2 isolates (Figure 1B). The sequences of the full-length complementary DNAs (cDNAs) of dsRNAs 1-5, were determined by assembling partial cDNA sequences amplified from each individually purified dsRNA using RT-PCR with tagged random primers and rapid amplification of cDNA ends (RACE). The assembled sequences of dsRNAs 1-5 were respectively 2399, 2188, 1965, 1131 and 1059 bp in size and were deposited in GenBank under accession numbers KT372135-KT372139. Following BLASTn searches of public databases, the dsRNAs sequenced in this study were found to be 99.0-100% identical to those of Botryosphaeria dothidea RNA virus 1 (BdRV1) dsRNAs 1-5 ^[7]^, constituting a strain of this virus. However, using the same experimental approach, we failed to determine the terminal sequences of any of the smaller RNAs, leading us to conclude that they lacked accessible free 3’ and 5’ termini and, consequently, might have a circular structure. We then obtained preparations enriched in these RNAs from strain XA-3, as previously described for circular viroid RNAs (last lane in Figure 1B), and subjected them to two-dimensional polyacrylamide gel electrophoresis (2D-PAGE) ^[9]^. Several RNA bands were detected in the second denaturing gel, which migrated slower than host RNAs (Figure 1C indicated by red arrows). Such behavior is consistent with the presence of circular RNAs ^[9]^. By contrast, the BdRV1 dsRNAs migrated together with host nucleic acid bands (Figure 1C, indicated by green arrows). These results suggest that the six small RNA bands visualized on gels are most likely circular RNAs and were accordingly nominated Botryosphaeria dothidea circular RNAs (BdcRNAs).

### 2.2 Full length sequences and strand polarity of the BdcRNAs

To obtain full-length sequences of each BdcRNA, abutted primer pairs of opposite polarity were designed, based on the assembled contigs of partial cDNAs, and amplified using RT-PCR with tagged random primers from each individually purified BdcRNA. Six amplified cDNA bands, corresponding to RNAs *ca*. 100-500 nt long were obtained (with primer pairs F1/R1, Figure S1A). Furthermore, RT-PCR using additional adjacent primers (F2/R2) of opposite polarity based on the sequences obtained for BdcRNAs 1, 2.1 and 3.1, produced cDNA bands of the expected sizes, further supporting their circular nature (Figure S1A). Sequence analysis of the full-length cDNAs from the six BdcRNAs indicated that they were respectively 450, 429, 396, 374, 221 and 157 nt in length and were designated as BdcRNAs 1, 2 (2.1, 2.2 and 2.3) and 3 (3.1 and 3.2 based on their sequence similarity.).

Both strands of BdcRNAs 1, 2.1, and 3.1 were labelled with DIG by *in vitro* transcription of their corresponding full-length cDNAs and their specificity was determined without any cross-hybridization between the strands under the experimental conditions employed (see below). Using DIG-labelled antisense probes, the circular nature of BdcRNAs 1 and 2 was confirmed following 2D-PAGE analysis in combination with northern-blot hybridization (Figures 1D and E).

To confirm the infectivity of BdcRNAs independent of BdRV1 or other potential co-infected mycoviruses, BdcRNA bands co-migrating with 200-500 bp markers were eluted from agarose gels and used to transfect strain MAO-2. Nucleic acids were extracted from the protoplast-generated sub-isolates using silica spin columns to enrich dsRNA levels ^[10]^. These extracts, which did not contain any BdRV1 dsRNAs (Figure 1FI) were subjected to dot-blot hybridisation using a DIG-labelled BdcRNA2.1 antisense probe and 4/12 protoplast-generated sub-isolates gave positive results (Figure 1FII). Further sub-cultures up to six generations were analysed by northern blotting with the same probe, illustrating that the BdcRNA2 titre and pattern were similar to those of strain XA-3, but higher titre and additional RNA molecules (including dimeric and circular forms) were noted as compared to the original transfectants (Figure 1G). Moreover, these transfectants were further confirmed free of BdRV1 while presence of BdcRNA2 by RT-PCR identification (Fig. SB). These results strongly support the notion that BdcRNAs are infectious and can initiate replication following protoplast transfection.

The positive-sense strands of these BdcRNAs were defined as those that accumulated to a higher titre compared to their counterparts in XA-3 mycelia, as accessed by northern blot analysis with the corresponding probes (see below).

### 2.3 BdcRNAs are single-stranded and highly paired

The single-stranded (ss) and highly paired nature of the BdcRNAs was established following digestion with DNase I, S1 nuclease, RNase III and RNase R, using as a control, citrus exocortis viroid (CEVd) circular ssRNA (Figure S1BI) in combination with dot-blot hybridization using BdcRNA2.1 and CEVd antisense probes (Figure S1BII). BdRV1 dsRNAs, remnant genomic DNA, and ribosomal RNAs, co-extracted from strain XA-3, were included as additional controls (Figure S1CI). The results showed that the circular RNAs extracted from strain XA-3, together with CEVd RNA, resisted digestion by DNase I, S1 nuclease, RNase III and RNase R (Figure S1BII). In parallel, the controls were digested by the corresponding enzymes: BdRV1 dsRNAs by RNase III, genomic DNAs by DNase I and ribosomal RNAs by S1 nuclease and RNase R, but remained unaffected when treated with other enzymes (Figure S1BI). Notably, the hybridization signals generated by BdcRNAs from strain XA-3 and CEVd RNA were marginally decreased following treatment with S1 nuclease and RNase R, because unpaired nucleotides were digested but paired regions remained intact following treatment with S1 nuclease because the RNAs are circular and have partially paired structures, while the linear ssRNAs (e.g., the replicative intermediate of the circular RNAs) were degraded and the circular counterparts remained intact following treatment with RNase R (Figure S1CII).

### 2.4 BdcRNAs are phylogenetically related but share no detectable homology with other known RNAs

BLASTn searches revealed that BdcRNAs 1-3 did not share detectable similarities with any sequences deposited in the National Center for Biotechnology Information (NCBI) database or host genomic sequences ^[11]^. However significant sequence similarities between the BdcRNAs themselves were noted e.g., 25.6-32.5% (BdcRNAs 1 and 2 or 3), 32.6-68.9% (BdcRNAs 2 and 3), 66.7-97.2% (BdcRNAs 2.1 to 2.3), and 86.1% (BdcRNAs 3.1 and 3.2). More specifically, BdcRNAs 2.1 to 2.3 differ from each other in deletions of *ca*. 25-30 nt, and BdcRNAs 3.1 and 3.2 in a deletion of a *ca*. 100 nt (Figure 2A).

**Figure 2.**
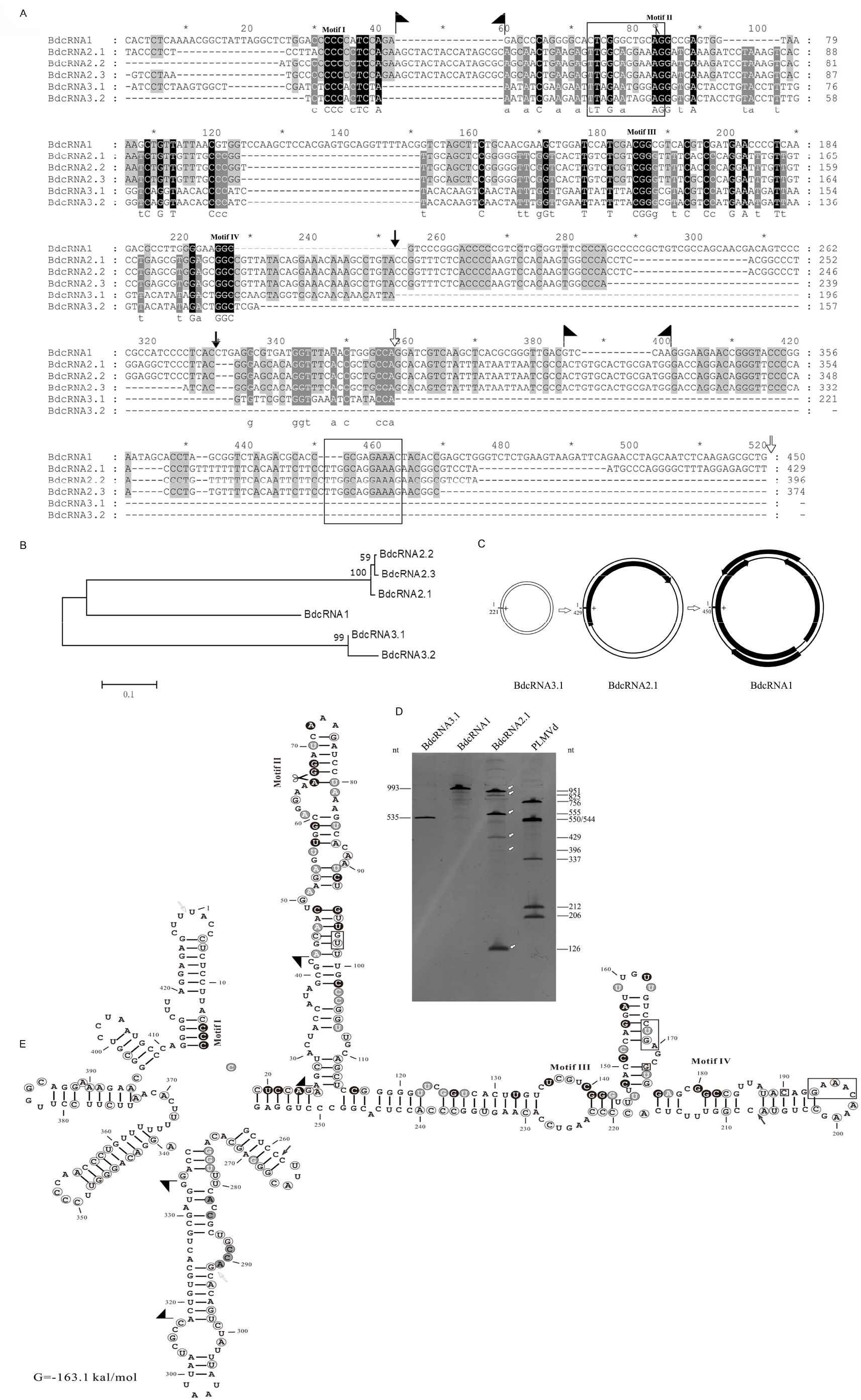
Sequence, phylogenetic, open reading frame (ORF), catalytic activity, and secondary structure analysis of BdcRNAs. (A) Multiple alignment of BdcRNAs using nucleotide acid sequences from MAFFT version 6.85 as implemented at http://www.ebi.ac.uk/Tools/msa/mafft/ with default settings, except for refinement with 10 iterations. (B) Phylogenetic relationships of BdcRNAs. (C) Schematic diagram of ORFs harbored by both strands of BdcRNAs. The double circular lines refer to the genome in plus (+) and minus (-), and black block refer to the ORFs with arrows indicating the orientation along the genome. (D) PAGE analysis of catalytic activity of transcription products derived from dimeric cDNAs of BdcRNAs and peach latent mosaic viroid (PLMVd). The numbers refer to the sizes of cleaved products of PLMVd dimeric RNAs or transcribed BdcRNA RNAs, and the arrows indicate the cleaved fragments. (E) The variation positions, conserved motifs and catalytic site indicated in the secondary structure of BdcRNA2.1. The secondary structures of BdcRNA1 to 3 as determined using the RNA Structure prediction tool in CLC RNA Workbench software (Version 4.8, CLC bio-A/S). Black, gray, and light gray backgrounds denote nucleotide identity conserved in all, five and four BdcRNAs, respectively. The arrows and flags denote deletion and insertion delimited positions, respectively, and the scissors denote self-cleavage site by the ribozyme.

Alignment of the BdcRNAs 1-3 sequences revealed four GC-rich motifs (I to IV) and a highly conserved 30 nt stretch (indicated with a black background in Figure 2A). Extensive sequence indels were also identified e.g., BdcRNAs 2 contain insertions (_43_AGCUACUACCAUAGCGC_59_ and _386_ACUGUGCACUGCGAUGG_399_, delimited by flags in Figure 2A) absent from the other BdcRNAs, while BdcRNA 3 lack *ca*. 190 nt corresponding to nt 253-329 and nt 357-520 of BdcRNA2.1 (indicated by arrows in Figure 2A). BdcRNAs 2 contains a 12 nt stretch (_73_UUGGCAGGAAAG_85_) which is directly repeated at nt 452-463, but is absent from BdcRNAs 3 while some remnants remain in BdcRNA1 as indicated by squares in Figure 2A.

BdcRNAs 1, 2.1, 2.2 and 2.3 have G+C contents of 59.3%, 53.9%, 54.3% and 53.7% respectively. By contrast, BdcRNAs 3.1 and 3.2 have lower G+C contents of 41.6 to 40.1%. Phylogenetic analysis revealed three clusters corresponding to BdcRNAs 1 to 3 (Figure 2B). The secondary structures of BdcRNAs 1 to 3, based on the lowest free energy predictions, show simple to complex branched and highly paired conformations differing in the number of loops and branches, respectively (Figures 2D and S2).

Open reading frame (ORF) prediction revealed that BdcRNA1 contains four ORFs, including one in the designated plus (+) strand and three in the minus (-) counterpart, potentially encoding small proteins with estimated molecular masses (*Mr*) s of 8.7, 3.2, 4.3 and 4.7 kDa, respectively (Figure 2C). A BLASTp search revealed that the largest putative protein has the highest identity (36.6%) with a hypothetical protein of unknown function (AS27_14953) from the emperor penguin *Aptenodytes forsteri* (KFM08303, coverage 51.0%, e-value 2.5). One small ORF was also detected in BdcRNA2, potentially encoding a protein with an estimated *Mr* of 6.8 kDa that shows highest identity (45.7%) with a hypothetical protein of unknown function (Saspl_004291) from the scarlet sage *Salvia splendens* (TEY57791, coverage 53.0%, e-value 4.1). No putative ORFs were detected in BdcRNA3 (Figure 2C). To check whether the putative proteins encoded by the BdcRNAs are *bona fide*, strains XA3-MAO-2-5 (harboring both BdRV1 and BdcRNA1), XA3-MAO-2-9 (harboring BdRV1 only) and MAO-2 were analyzed by LC-MS after trypsinization. No peptides matching these putative proteins in the aforementioned strains were found, while BdRV1 proteins including the RNA dependent polymerase (RdRp) and coat protein were found in both XA3-MAO-2-5 and XA3-MAO-2-9. Overall, these results suggest that the BdcRNAs ORFs most likely do not encode any proteins.

The potential self-catalytic activities of BdcRNAs were examined using RNA transcribed from recombinant plasmids containing dimeric inserts. Dimeric RNAs of the peach latent mosaic viroid (PLMVd) containing a HHRz were used as a control. BdcRNA2.1 showed self-catalytic activity resulting in individual RNA products 941, 823, 547, 429, 394, and 118 nt in size, in contrast to both BdcRNA1 and BdcRNA3.1 (Figure 2D). RACE, cloning and sequencing of the smallest cleaved RNA products of BdcRNA2.1 revealed the catalytic site between nt A_66_ and G_67_ (Figure 2 E), within motif II which is conserved in all BdcRNAs (Figure 2A). Notably, BdcRNA2.1 is a novel ribozyme, which is distinct from all known ribozymes including viroid hammerhead ribozymes.

### 2.5 BdcRNA(s) are horizontally and vertically transmitted independently

Horizontal transmissibility of BdcRNAs was addressed by contact cultures between XA-3 (donor) and MAO-1 or MAO-2 (recipient) strains of *B. dothidea* in different combinations (Figure S3AI and II). The colony morphology of several sub-isolates derived from contact cultures was monitored, together with the presence of RNA elements, using dsRNA extraction for BdRV1 and dot-blot hybridization for BdcRNAs (with antisense probes). Most sub-isolates displayed colony features similar to those of their parental strains (i.e., MAO-1 sub-isolates were dense and cotton thread-like, while MAO-2 sub-isolates were thin and sparse in the centre but some were highly divergent (e.g., MAO-1-3 and −7, and MAO-2-5) (Figure S3AIII). The observed changes were not due to contamination, as confirmed by sequencing the ITS regions of the divergent sub-isolates. Seven out of nine MAO-2 sub-isolates (MAO-2-1 to −7) were infected only by two or three BdcRNAs (MAO-2-1, −4 and −5), in some cases together with BdRV1 (MAO-2-2, −3, −6 and −7); seven out of ten MAO-1 sub-isolates were infected by only one to three BdcRNA(s) (MAO-1-5 and −7), in some cases with BdRV1 (MAO-1-3, −6, −8, −9 and −10) (Figure S3B).

To investigate the vertical transmissibility of BdcRNAs, XA-3 conidia were isolated by inoculating mycelium discs on young fruits for sporulation. Examination of BdcRNAs by dot-blot hybridization (with antisense probes) and of BdRV1 dsRNA patterns in 20 randomly chosen sub-isolates of conidium cultures showed that all were infected by BdcRNAs1 to 3 and by BdRV1 (Figure S3C), thus supporting the efficient transmission of BdcRNAs through *B. dothidea* conidia.

### 2.6 BdcRNA(s) have different subcellular localizations and variable self-catalytic activities

To gain insight as to the subcellular locations of the BdcRNAs, fluorescence *in situ* hybridization (FISH) was performed with XA-3 protoplasts using Alexa Fluor 488-labeled riboprobes specific for (+) and (-) strands of BdcRNA1 and BdcRNA2.1. Both strands of BdcRNA1 were present predominantly in the nuclear peripheral regions (Figure 3AI), whereas these of BdcRNA2 were concentrated in nuclei (Figure 3AII), as shown by green fluorescence produced by the probes overlapping red fluorescence produced by mCherry, which binds to fungal histones, and blue coloration following staining with diamidine phenylindole (DAPI) (Figure 3A). Conversely, in some cells both strands were absent from nuclei but still diffused into the cytoplasm (Figure S4A and B). In mycelia, both strands of BdcRNA1 and the (+) strands of BdcRNA2 but not the (-) strands systemically and uniformly spread in growing or newly formed mycelia where nuclei were yet to mature (Figure 3B). No hybridization signals were found in MAO-2 cells which do not harbor BdcRNAs (Figure 3C).

**Figure 3.**
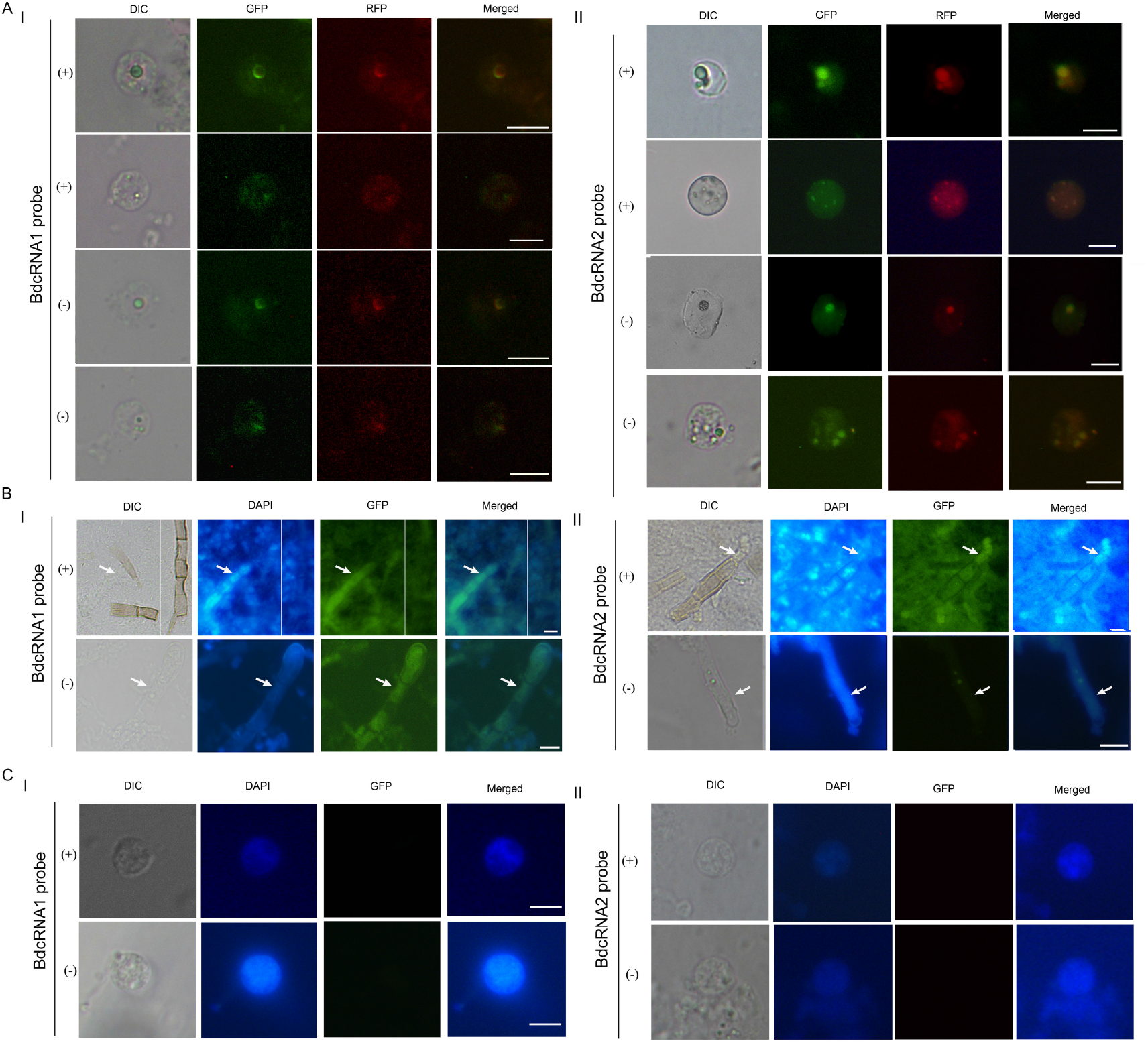
Subcellular location of BdcRNAs. (A and B) Fluorescence *in situ* hybridization (FISH) to detect subcellular location of (+) and (-) strands of BdcRNA1 (I) and BdcRNA2.1 (II) in protoplasts (A) of strain XA-3, by Alexa Fluor 488-labelled riboprobes (green fluorescence), as well as in (B) newly formed mycelia containing no assembled nuclei, with strain MAO-2 protoplasts investigated in parallel as controls (C). The representative protoplasts containing one and more nuclei are presented as observed under differential interference contrast (DIC) or fluorescent settings. Nuclear location was indicated with mCherry, binding to the fungal nucleosome (red fluorescence), or stained with DAPI blue color at a higher concentration than in the cytoplasm. The fungal cells contain none or one or more nuclei depending on their age and those nuclei in the process of assembling are unlabeled by mCherry or stained with DAPI at a high intensity.

### 2.7 BdcRNA (s) replicate(s) *via* a rolling-circle mechanism following symmetric pathways *in vivo*

DIG-labeled full-length sense and antisense specific probes of BdcRNAs 1, 2.1, and 3.1 detected anti-polarity transcripts (Figure 4A, lanes 2, 10, 15, 23, 29, and 37), but did not cross hybridize with sense-polarity transcripts (lanes 3, 9, 16, 22, 30, and 36) and nucleic acids from MAO-2 (lanes 1, 8, 21, 28, 35, and 42). Northern hybridization of RNA preparations from XA-3 with similar amounts of these probes revealed that the (+) ssRNAs of BdcRNAs 1 to 3 accumulated abundantly *in vivo* in circular forms resistant to RNase R digestion (Figure 4A, lanes 5, 18, and 32), together with their dimeric (for BdcRNAs 1 and 2; lanes 4 and 17) or trimeric forms (for BdcRNA3; lane 31) in substantially lower titres as compared to the circular forms. The (-) ssRNAs of BdcRNAs 1 to 3 also accumulated in circular forms (Figure 4A, lanes 12, 25, and 39) together with the dimeric (for BdcRNAs 1 and 2; lanes 11 and 24) or oligomeric forms (for BdcRNA3; lane 38). These results suggest that BdcRNAs may adopt a rolling-circle replication mechanism following a symmetric pathway, similar to that adopted by chloroplastic viroids. Here a circular (+) strand is transcribed to yield a dimeric (for BdcRNAs 1 and 2) or an oligomeric (for BdcRNA3) linear (-) strand, which is cleaved by host protein(s) (for BdcRNAs 1 and 3; Figure 4BI) or by the ribozyme (for BdcRNA2; Figure 4BII) to unit lengths and then circularized; the resulting (-) circular strand serves as a template to produce (+) dimeric (for BdcRNAs 1 and 2) or oligomeric forms (for BdcRNA3), which are cleaved and then circularized in the nucleus. The resulted circular (+) strands are transported into the cytoplasm (Figure 4B).

**Figure 4.**
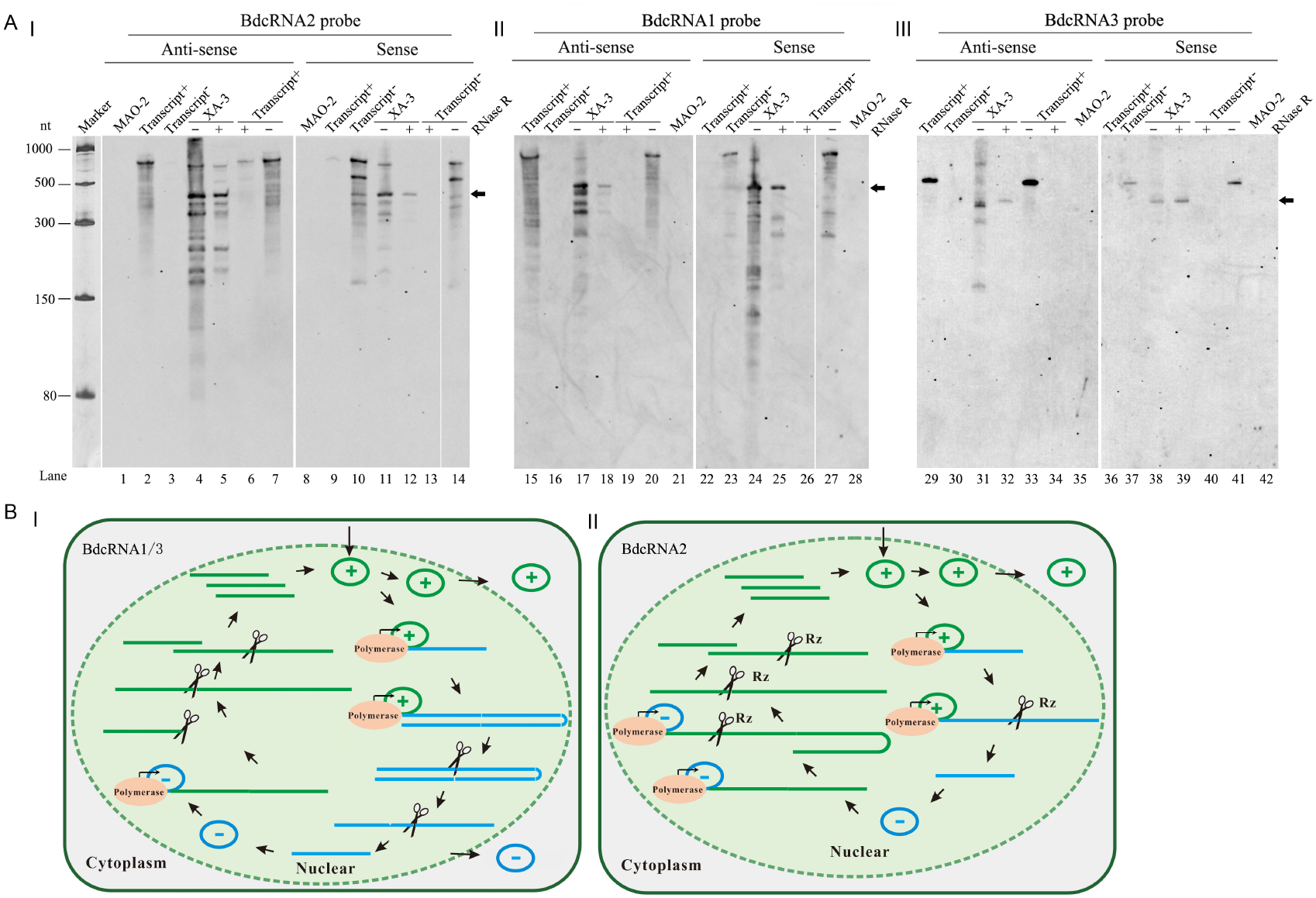
Replication analysis of BdcRNAs. (A) Northern blot hybridization of nucleic acid preparations from strains MAO-2 and XA-3 using antisense and sense probes of BdcRNAs 1 (II), 2.1 (I) and 3.1 (III), respectively. Marker 1, an ssRNA marker electrophoresed on the same gel was visualized by staining with silver nitrate. The (+) and (-) ssRNAs of BdcRNAs derived from *in vitro* transcription were examined to assess probe specificity and confirm lack of cross hybridization between strands (lanes 2, 3, 9, 10, 15, 16, 22, 23, 29, 30, 36, and 37). The nucleic acid preparations from strain XA-3 were also subjected to digestion with RNase R to confirm their circular nature, together with the transcripts binding to the probes as controls. In this case “+” and “-” refer to presence and absence of RNase R, respectively. Lanes are numbered below the panels. The arrows indicate circular forms. (B) Proposed symmetric models of rolling-replication pathway for BdcRNAs 1 and 3 (I), and 2 (II), respectively.

### 2.8 Artificial BdcRNA(s) replicate autonomously and spread systemically *in vivo*

To assess the infectivity and replication of BdcRNAs, *in vitro* transcripts derived from dimeric head-to-tail cDNAs of BdcRNA1, 2.1 and 3.1 were transfected into MAO-2 protoplasts resulting in respectively thirteen, four and eleven positive sub-isolates, out of 50 as assessed by dot-blot hybridization and RT-PCR amplification (Figure S5).

To determine whether BdcRNAs can spread systemically following protoplast transfection, three transfectants (T11, T32, and T14 transfected with BdcRNAs 1, 2.1 and 3.1, respectively) were selected for protoplast isolation and twenty single cells, designated T11-1 to −20 (for BdcRNA1), T32-1 to −20 (for BdcRNA2.1), and T14-1 to −20 (for BdcRNA3.1), were serially cultured over six generations. The BdcRNA titres of the transfectants, together with their parent strains (T11, T32, and T14), were assessed by RT-qPCR amplification (flowchart, Figure S6A). Based on the quantitative absolute standard curves established with serially diluted plasmids containing BdcRNA cDNAs (Figure S6B), RT-qPCR amplification revealed that BdcRNA1 titers fluctuate (1.2-1.7×10^4^ copies μL^−1^) in the progeny (T11-1 to −20) without any significant difference as compared to the titre (1.5 × 10^4^ copies.μL^−1^) of the parent isolate (T11) (Figure S6CI). The results for BdcRNA2.1 and BdcRNA3.1 (T32-1 to-20 and T14-1 to −20) were similar, but fluctuations were much more prominent and progeny with substantially higher or lower titers were recorded (Figure S6C II and III). These results suggest that BdcRNAs can replicate autonomously and spread systemically into newly growing cells, but that their accumulated titers differ between individual-cell generated colonies.

### 2.9 BdcRNAs accumulation is not affected by co-infection with mycovirus

To determine whether replication of BdcRNAs is affected by BdRV1 co-infection, BdRV1 was introduced to sub-isolate T11 using horizontal transmission, resulting in three positive sub-isolates (BdRV1-T11-C5 to −C7) as ten were identified by dsRNA extraction (Figure S7AI). The BdcRNA1 titres of three resulting progenies infected by BdRV1 were assessed by qRT-PCR, revealing fluctuating titres in the BdRV1-infected sub-isolates but with no significant difference as compared to the parent sub-isolates (Figure S7AII).

Furthermore, to determine whether replication of BdcRNAs is affected by other viruses, Botryosphaeria dothidea partitivirus 1 (BdPV1) was introduced into subisolates T11 (BdcRNA1^+^) and T32 (BdcRNA2.1^+^) using protoplast transfection and then examined by dsRNA extraction (Figure S7B). Ten transfectants from each sub-isolate were assessed by qRT-PCR, revealing titres in the BdPV1-infected sub-isolates similar to those of the parent sub-isolates (Figure S7CI and II). These results suggest that mycovirus co-infection had no obvious effects on the accumulation of BdRNAs *in vivo*.

### 2.10 Individual BdcRNA(s) modulate specific biological traits of the host fungus

To better understand the biological roles of individual BdcRNAs, six of each derived transfectants of BdcRNA 1, 2.1 and 3.1, together with XA-3, MAO-1, and MAO-2, were grown on PDA and complete medium (CM). BdcRNA1 and BdcRNA2.1 transfectants (T11-1 to −10, and T32-1 to −17) had altered morphology as compared to the parent strain MAO-2: the former produced dense mycelia, while the latter produced sparse mycelia with irregular margins; in contrast, BdcRNA3.1 transfectants (T14-3 to −53) showed no obvious changes (Figure 5A). Moreover, the growth rates were significantly decreased for both BdcRNA1 and BdcRNA2.1 transfectants but not for BdcRNA3.1 transfectants (Figure 5B). Additionally, BdcRNA1 virtually eliminated fungal virulence as assessed on pear fruits and branches, since no lesions were observed following transfectant infection, while the parent strain MAO-2 resulted in lesions *ca*. 18.1 mm in size on fruits and *ca*. 20.7 mm on branches; BdcRNA2.1 transfectants also exhibited significantly decreased virulence, whereas the BdcRNA3.1 ones did not (Figures 5C and D). These results suggest that BdcRNAs play diverse but well-defined roles in the modulation of host morphology, growth, and virulence.

**Figure 5.**
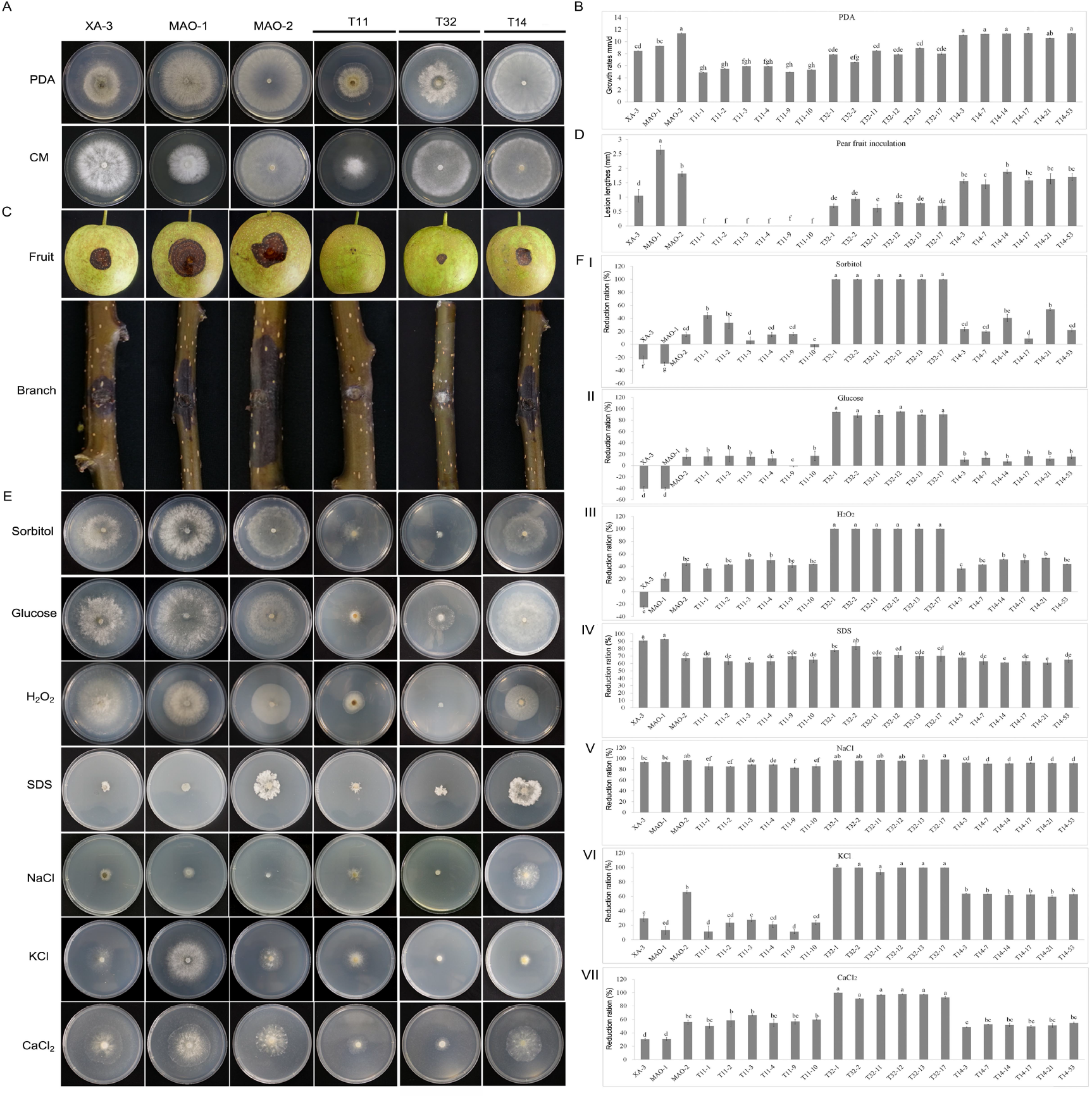
Biological traits of transfectants of *B. dothidea* MAO-2 infected by BdcRNAs together with control strains. (A and B) The representative morphologies of the transfectants derived from BdcRNAs 1 (T11-), 2.1 (T32-) and 3.1 (T14-) on PDA and complete medium (CM), and the histogram for their growth rates assessment on PDA (n=3), respectively. (C and D) Virulence assessment on pear (var. Cuiguan) fruits (n=8) and tree branches (n=6) for seven days, and the histogram for the resulting lesions, respectively. (E and F) Growth assessment on PDA amended with 1 mol L^−1^ sorbitol (I), 1 mol L^−1^ glucose (II), 10mM H2O2 (III), 0.005% SDS (IV), 1.5 mol L^−1^ NaCl (V), 1 mol L^−1^ KCl (VI), and 0.5 mol L^−1^ CaCl2 (VII) incubated at 25°C in darkness for three days, and the histograms of growth reduction rates as compared with the growth on PDA (n=3), respectively. Bars in each histogram labeled with the same letters are not significantly different (P >0.05) according to the least significant difference test (1-way ANOVA); and error bars represent SD.

To determine the role of BdcRNAs in fungal stress responses, transfectants were cultured on PDA amended with different cellular stress agents, including those involved in sugar metabolism (1 mol L^−1^ sorbitol and 1 mol L^−1^ glucose), cell wall disruption (0.005% SDS), oxidative stress (10 mM H_2_O_2_), osmotic stress (1 mol L^−1^ KCl and 1.5 mol L^−1^ NaCl), and the calcium pathway (0.5 mol L^−1^ CaCl_2_). All BdcRNA1 transfectants (T11-1 to −10) showed significantly higher tolerance to high concentrations of KCl and NaCl as compared to the parent strain MAO-2; BdcRNA2 transfectants showed strong sensitivity to high concentrations of sorbitol, glucose, H_2_O_2_, KCl, and CaCl_2_; BdcRNA3 transfectants (T14-3 to −53) only showed tolerance to high concentration of NaCl (Figure 5E and F). These results showed that individual BdcRNAs interfere with functions of *B. dothidea* strains and exert diverse regulatory roles *in vivo* under different stress conditions.

### 2.11 BdcRNA(s) modulate gene expression and metabolism of the host fungus

To analyze the effect of BdcRNAs on host gene expression, three transfectants together with the parental strain of each BdcRNA were subjected to transcriptome profiling by next-generation sequencing and differential gene expression analysis following culture on PDA for 8 days. The results revealed distinct gene expression patterns modulated by BdcRNAs 1 to 3 as compared to the control strain MAO-2 according to heat map analysis (Figure 6A), with 6741, 2122 and 3400 differentially expressed genes (DEGs) (Figure 6B), accounting respectively for 47.8%, 15% and 24.1% of the total genes of the host fungus (14116 genes in the genome, ID: 12428). Gene ontology (GO) classification revealed the DEGs were mainly involved with peptidase metabolic pathways, including peptidase, endopeptidase, serine-type endopeptidase, and serine-type peptidase activities for BdcRNA1 transfectants (Figure 6CI), ribosome metabolic pathways (rRNA metabolic process, rRNA processing, ribosome biogenesis, and ribonucleoprotein complex biogenesis) for BdcRNA2 transfectants (Figure 6CII), and proteolysis, signaling (signaling, signal transduction, single organism signaling), and cell communication for BdcRNA3 transfectants (Figure 6CIII).

**Figure 6.**
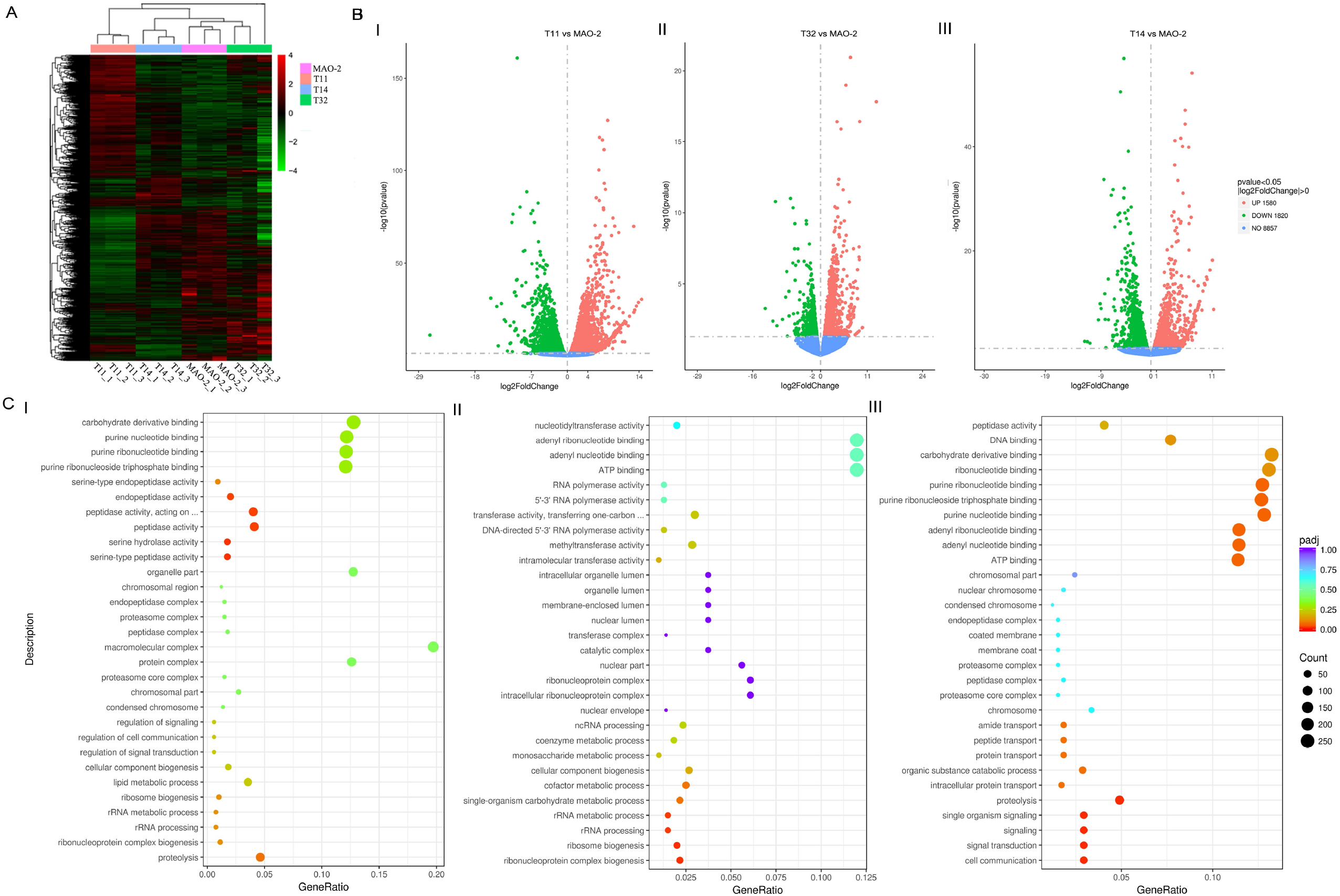
Transcriptome analyses of transfectants of *B. dothidea* MAO-2 infected by BdcRNAs together with the control strain. (A) Clustered heat map of differentially expressed genes (DEG) of the transfectants infected by BdcRNAs 1 (T11), 2.1 (T32) and 3.1 (T14) together with MAO-2. (B) Volcano plots of DEG of transfectants T11, T32 and T14 as compared with MAO-2. (C) Scatter diagrams of GO enrichment analysis of DEG of transfectants T11 (I), T32 (II) and T14 (III).

To verify the transcriptome results, six downregulated DEGs, including these of the glycosyl hydrolase family (ID A08082), the glycosyl transferase family (ID A03050) and some of unknown functions (A07378, A06265, A03482 and A09837), were examined by RT-qPCR in MAO-2 transfectants (T11 and T32) at 6, 8 and 10 dpi on PDA. All selected genes showed decreased expression patterns similar to those obtained by transcriptome profiling at no less than one time point, supporting the next generation sequencing outcome (Figure S8).

## 3. Discussion

In this study we have identified a set of circular ssRNAs, BdcRNAs-1 to −3, in *B. dothidea* isolate XA-3. Understanding the nature of these molecules was based on the following evidence: (i) 2D-PAGE in combination with northern-blot hybridization revealed that BdcRNAs migrated slower than host RNAs of comparable size, (ii) BdcRNAs were resistant to DNase I, RNase R and RNase III, and (iii) RT-PCR amplification using abutted primer pairs amplified full-length sequences. BdcRNA(s) were capable of horizontal transmission, independent of each other or mycoviruses, and autonomous replication following transfection.

BdcRNAs display a rapid genomic expansion including insertions, mutations, and deletions, and they show remarkable variation in nucleic acid sequence, size (175-450 nt), secondary structure complexity, GC content and protein coding capacity. However, BdcRNAs share nucleotide identities including four conserved GC-rich motifs and a highly conserved stretch >30 nt in length (Figure 2A). These observation support evolution of BdcRNAs from a common ancestor. Notably, BdcRNA1 and 2 contain potential ORFs, which clearly separate them from BdcRNA3 (and viroids); however, no resulting proteins were detected by proteomics analysis, which might provide a link between noncoding and coding RNAs. Since the properties of BdcRNA(s) are those expected for primitive RNA replicons, including small size (BdcRNA3.1), high GC content (up to 59.3% for BdcRNA1), circular structure, structural periodicity, lack of coding capacity (for BdcRNA3), and a novel ribozyme (for BdcRNA2), i.e., the fingerprints of the RNA world ^[1a]^, BdcRNA(s) may represent a link between primordial RNAs and modern coding RNAs.

BdcRNAs share no detectable sequence identities with any of these endogenous cRNAs, or with the host genome, and are not detectable in virulent *B. dothidea* strains, suggesting that they are exogenous cRNAs. The known exogenous small cRNAs in eukaryotic cells include small satellite RNAs and viroids. Satellite RNAs have genomes *ca*. 350 nt in length, infect plants and are associated with plant viruses. Two cherry small circular RNAs (cscRNA1 and cscRNA2) have been isolated from cherry trees, are found in combination with mycoviral dsRNAs and it has been suggested that they are viroid-like satellite RNAs ^[12]^. BdcRNA(s) are independently infectious, and their replication is unaffected by mycoviruses, suggesting that they are not satellite RNAs or viroid-like satellite RNAs but another type of infectious agent resembling viroids in their autonomous replication. In particular, BdcRNAs resemble viroids infecting plants in terms of size, circularity, and infectivity, but they lack the CCR and HHRzs typical of nuclear and chloroplastic viroids, respectively. Additionally, BdcRNAs have strands located in both the nucleus and the cytoplasm with distinct distribution patterns, significantly different from the subcellular location of *Pospiviroidae* and *Avsunviroidae* viroids, which are located in the nucleus and the chloroplast, respectively, with both of their strands (as demonstrated in CEVd, coconut cadang cadang viroid, and PSTVd) exhibiting similar distribution patterns ^[13]^. Furthermore, although BdcRNAs 1 and 3 contain complex branched secondary structures similar to those of chloroplastic viroids, they possess no ribozyme similar to nuclear viroids ^[1b, 14]^. BdcRNA2 contains a novel ribozyme, which is distinct from other ribozymes including the HHRzs. Therefore, BdcRNAs may represent a novel class of autonomously replicating subviral agents which are not satellites but resemble plant viroids. In previous studies, avocado sunblotch viroid (ASBVd) was shown capable of infecting the unicellular fungus *Saccharomyces cerevisiae* with their dimeric/oligomeric cDNAs fused in an expression vector ^[3a]^; ASBVd, hop stunt viroid (HSVd), and iresine viroid-1 (IrVd-1) were respectively transfected with monomeric transcripts into three filamentous phytopathogenic fungi (*Cryphonectria parasitica, Valsa mali*, and *Fusarium graminearum*) These results support the notion that viroids could replicate in at least one of these three fungi ^[3e, 15]^, although this study has stimulated some controversy since the major evidence for their replication in the fungi was derived by RT-PCR, and a reassessment (with a second detection technique, appropriate controls, and further experiments) is needed to support claims for infectivity in fungi ^[3e, 16]^. No natural infections of fungi with viroids or viroid-like RNAs have been discovered in previous studies ^[17]^ and, to our knowledge, this is the first report of infectious viroid-like RNAs (or exogenous small circular RNAs) in a life kingdom (fungi) other than plants. Here, the term ‘mycoviroid’ was tentatively used for such types of novel acellular entities with reference to viroid-like RNAs naturally infecting fungi, expanding the term beyond its original definition as “a viroid that has the ability to infect healthy fungi” ^[18]^.

FISH experiments revealed that BdcRNAs have different distribution patterns in both cells and mycelia depending on the species under investigation and RNA polarity. High concentrations of both strands in the nucleus may reflect their localization during replication and transcription. The absence of both strands of BdcRNAs 1 and 2 from nuclei observed in some cells is likely caused by a cessation of replication due to cell aging and the export of synthesized cRNAs out of the nuclei. Correspondingly, BdcRNA2 (+) strands instead of (-) strands were translocated from older to younger hyphae. Detection of the (+) and (-) dimeric or oligomeric forms together with circulars form in both polarities suggests that BdcRNAs 1 to 3 follow a symmetric rolling-circle replication mechanism *in vivo* (Figure 4B), similar to previous reports for viroids ^[19]^. The absence of oligomeric forms for BdcRNAs 1 and 2, unlike the avocado sunblotch viroid, which replicates using a symmetric replication pathway ^[20]^, might be due to rapid self-cleavage during elongation. To date, no information concerning the mechanisms of replication and movement of exogenous cRNA in fungi have been reported. Here we demonstrate for the first time that exogenous small cRNAs (in diverse species) replicate, are transported, and distributed in fungal cells in diverse ways depending on the species and strand polarity. These observations support the notion that fungi have developed machinery which can recognize and localize diverse types of small cRNAs in a strand specific manner.

BdcRNAs significantly affect the biological traits of *B. dothidea*, e.g., alter morphology, decrease growth rate, attenuate virulence, and increase or decrease tolerance to osmotic stress and oxidative stress. Concurrent with phenotype modulation, gene expression and metabolic pathways related to important cellular processes of the fungal host were also modulated by BdcRNAs. For BdcRNA1 and 2-transfectants, all related RNA processing genes were significantly downregulated and these changes may provide an explanation as to why BdcRNAs-infected strains have attenuated growth rates. RNA processing is important in biology, involving nucleic acid metabolism, gene expression, protein metabolism and cellular component biogenesis ^[21]^, all of which play crucial roles in fungi. Moreover, the DEGs related to peptidase activities were significantly downregulated for BdcRNA1 transfectants. These observations may explain their attenuated virulence since peptidases (proteases) contribute to both fungal growth and pathogenesis, and are considered markers of pathogenicity ^[22]^. Virulence was differentially attenuated in all BdcRNA transfectants, most likely due to the extent of differential expression of related genes. Thus far, mycoviruses are the only acellular agents that have been extensively investigated and infect all major fungal taxa. Here we describe another acellular entity apart from mycoviruses which confer important effects on the physiology and pathogenicity of the host fungus, as exemplified by BdcRNA1 that dramatically reduces host virulence. This feature provides an important alternative candidate to serve as a biocontrol tool for attenuation of fungal diseases similar to some mycoviruses that cause hypovirulence ^[23]^.

In summary, this work shows that a set of BdcRNAs display molecular and biological features unreported in known living entities, and may represent a new class of viroid-like RNAs endowed with regulatory functions, and novel epigenomic carriers of biological information. BdcRNAs contribute to the diversity of fungal biological traits, and may help us understand fungal diversity and biological changes along the life process besides genomics of fungi or even other cellular organisms. Moreover, BdcRNAs can be regarded as additional inhabitants of the frontier of life in terms of genomic complexity, and may represent another group of “living fossils” from a precellular RNA world as compared with viroids ^[1a, 24]^. More importantly, BdcRNAs can attenuate the virulence of phytopathogenic fungi, and may be exploited as biocontrol agents. Here, “mycoviroid” was termed for such viroid-like RNAs naturally infecting fungi, with BdcRNAs 1 and 2 as the type species of two separate groups.

## 4. Experimental Section

### 4.1 Fungal isolates and biological characterization

*B. dothidea* strains XA-3, MAO-1 and MAO-2 were isolated from apple tree branches (*Malus domestica* Borkh. cv. ‘Fuji’) collected in Shandong province, China and identified based on morphological characteristics and molecular analyses. All strains were purified by hyphal-tipping ^[25]^.

### 4.2 RNA extraction and enzymatic treatments

For dsRNA extraction, mycelial plugs were inoculated onto cellophane membranes on PDA (20% diced potatoes, 2% glucose and 1.5% agar) plates and incubated in darkness at 25°C for 4 to 5 days. Approximately 1 g mycelia were collected, ground to a fine powder in liquid nitrogen, and subjected to dsRNA extraction as previously described ^[10]^.Briefly, the frozen powder was treated with five volumes of buffer (2% SDS, 4% PVP-40, 0.5 M NaCl, 100 mM Tris-HCl (pH 8.0), 20 mM EDTA), and an equal volume of binding buffer (50% guanidine thiocyanate, 1.5 M KCl, 0.5 M NH_4_Cl, 0.3 M KAcO pH 6.0). The resulting mixture was centrifuged at 12,000 × g for 5 min. The supernatant was mixed with 0.5 volumes of ethanol, and loaded onto a silica spin column (Sangon Biotech (Shanghai) Co., Ltd, China) to absorb the dsRNA, which was eluted with 60 μL RNase-free water after two washes with the binding buffer containing 37% ethanol.

Alternatively, the frozen powder was subjected to extraction of RNAs with compact secondary structure as previously described ^[8]^. Briefly, frozen powder from 30 g mycelia was treated with buffer-saturated phenol (pH 7.0), and following centrifugation at 12,000 × g the supernatant was mixed with non-ionic cellulose (CF-11, Whatman) in 1×STE buffer (50 mM Tris-HC1 (pH 7.2), 100 mM NaC1 and 1 mM EDTA) and 30% ethanol and shaken overnight. After washing the cellulose three times with STE containing 35% ethanol, the bound RNAs (and other accompanying substances) were eluted with 1×STE, precipitated with ethanol, dissolved in 60 μL RNase-free water, and stored at −70 °C until use.

The RNA preparation from XA-3 together with circular and linear CEVd RNAs, extracted from a citrus tree infected by CEVd variant CEVd.188, ^[26]^ were subsequently treated with several enzymes to determine their nucleic acid nature. Briefly, aliquots of 200 ng nucleic acid were treated with 2 U DNase I (New England Biolabs), 10 U S1 nuclease (Thermo Scientific), or 0.4 U RNase III (New England Biolabs) in cleavage buffer [30 mM Tris (pH 8.0), 160 mM NaCl, 0.1 mM EDTA, 0.1 mM DTT, 10 mM MgCl_2_], as previously described ^[10]^, and with 200 ng mL^−1^ RNase R (Thermo Scientific) in 2×SSC (300 mM NaCl, 30 mM sodium citrate, pH 7.0) at 37°C for 1 h according to the manufacturer’s instructions. After enzyme treatment nucleic acids were treated with phenol/chloroform/isoamyl alcohol (25:24:1) (phenol saturated with water, pH 5.2), precipitated with ethanol, dissolved in RNase-free water, and stored at −70 °C until use.

The nucleic acid preparations were analyzed by 6% non-denaturing PAGE or 1.2% agarose gel electrophoresis and visualized by staining with silver nitrate and ethidium bromide, respectively.

### 4.3 cDNA synthesis and molecular cloning

The cDNA sequences of the five dsRNAs identified on gels were determined as previously described ^[27]^. The 5’ and 3’ terminal sequences of the dsRNAs were determined using RNA ligase mediated (RLM) RACE ^[27]^.

The circular RNA cDNA sequences were determined using a random cloning method as previously reported ^[10]^. Briefly, purified cRNAs were subjected to cDNA synthesis using M-MLV reverse transcriptase with Primer I (5’-GAC GTC CAG ATC GCG ATT TCN NNN NN-3’), and amplified using Primer II (5 ‘-GAC GTC CAG ATC GCG ATT TC-3’) in combination with end filling with *Taq* polymerase. The amplified PCR products were cloned into the pMD18-T vector (TaKaRa, Dalian, China) and transformed into competent cells of *Escherichia coli* DH5alfa. At least three independent clones of each fragment were sequenced in both directions. Sequencing was performed at Sangon Biotech (Shanghai) Co., Ltd, China, and each nucleotide was determined in at least three independent overlapping clones in both orientations.

Further RT-PCR identification of circular RNAs was conducted using the primers designed based on assembled contigs of cDNAs synthesized using random hexamers as described above. Amplification was performed in a thermal cycler for 35 cycles of 94°C for 30 s, 50-60°C for 30 s and 72°C for 30 s after initial denaturation at 95°C for 30 s, and followed by a final extension at 72°C for 10 min.

### 4.4 Sequence analysis

Sequence similarity searches were performed using the NCBI databases with the BLAST program. Multiple alignments of nucleotide and amino acid sequences were conducted using MAFFT version 6.85 as implemented at http://www.ebi.ac.uk/Tools/msa/mafft/ with the default settings except for refinement with 10 iterations. Identity analyses were conducted using the DNAMAN DNA analysis software package (DNAMAN version 6.0; Lynnon Biosoft, Montreal, Canada). The secondary structures of BdcRNAs were determined using the RNA structure prediction tool in CLC RNA Workbench software (Version 4.8, CLC bio A/S) with the algorithms as described ^[28]^. Open reading frames (ORFs) were deduced using ORF Finder online (https://www.ncbi.nlm.nih.gov/orffinder/).

### 4.5 2D-PAGE, probe preparation, and hybridization

2D-PAGE was conducted as previously described ^[8–9]^, with some modifications. Briefly, aliquots were first separated by non-denaturing PAGE in a 5% gel in 1×TAE (40 mM Tris-HC1, 20 mM NaAcO, 2 mM EDTA, pH 7.5) that was stained with ethidium bromide. The segment delimited by the DNA markers of 100 and 500 bp was excised and placed on top of a second denaturing 5% gel in 0.25×TBE (where 1×TBE buffer is 89 mM Tris, 89 mM H_3_BO_3_, 20 mM EDTA, pH 8.0) and 8 M urea, in which circular RNAs display slower electrophoretic mobility than their linear counterparts ^[8–9]^.

DIG-labeled full-length riboprobes complementary (antisense) or homologous (sense) to BdcRNAs 1, 2.1, and 3.1 were prepared as previously described ^[8, 29]^. Nucleic acids were spotted (for dot-blot) or electro transferred (for northern-blot) to positively-charged nylon membranes (Roche Diagnostics), and hybridized with DIG-labeled full-length riboprobes as previously described ^[8, 29]^.

### 4.6 Fluorescence *in situ* hybridization

FISH was conducted essentially as described ^[13]^, using probes labeled with Alexa Fluor 488 following the approach described below for DIG-labelled probes. All samples were stained with the DNA fluorochrome 4’,6-diamidino-2-phenylindole dihydrochloride (DAPI) dissolved in PBS at a final concentration of 1 μg mL^−1^ for 2 min. Additionally, strains MAO-2 and XA-3 were transfected with the pBS-NEO-Histone-mCherry plasmid and mCherry was used as an indicator of fungal histones in the nucleus. The excitation and emission wavelengths were respectively 330 to 380 for DAPI, 450 to 490 nm for Alexa Fluor 488, and 587 nm and 610 nm for mCherry.

### 4.7 Preparation of protoplasts and transfection of BdcRNAs

Protoplasts were prepared from actively growing mycelia of the corresponding strains as previously described ^[30]^ and aliquots stored at −70°C until use.

Protoplasts were filtered through a Millipore filter, counted under a microscope (×100) using a hemocytometer, and used for BdcRNA transfection -using PEG 6000 as previously described ^[31]^. A total of 1.0×10^6^ protoplasts were transfected with ~5.0 μg of *in vitro* transcribed dimeric BdcRNAs RNAs. Following transfection, the protoplast suspensions were diluted with sterilized water and spread onto PDA plates and any new colonies were separately cultured on fresh PDA plates for BdcRNA extraction.

### 4.8 Morphology, growth rate, stress-resistance, and virulence assays

Morphology and growth rate were estimated in triplicate at 25 °C following incubation in darkness at 3 dpi as previously described ^[10]^. The stress-resistance assay to salt and osmotic pressure was conducted by assessment of the growth rates at 25°C in the dark on PDA by supplying with corresponding components in suitable concentration to the medium after assessment of serial dilutions. The virulence of each strain was determined by inoculating detached fruits (in octuplicate) or branches (in sextuplicate) of *M. domestica* cv. ‘Fuji’ or *Pyrus pyrifolia* cv. ‘Cuiguan’ as previously described ^[32]^.

### 4.9 Quantitative real-time PCR

Quantitative real-time PCR (qRT-PCR) to measure BdcRNA titer was conducted using the CFX96 Real-Time PCR Detection System (Bio-Rad, USA) based on the cDNAs synthesized as in the - description above using specific primers designed for quantitative analysis. Amplification was conducted in a thermal circular for 45 cycles of 95°C for 30 s, 60 °C for 20 s and 72°C for 20 s after initial heating at 95°C for 3 min, and followed by a final extension for 10 min at 72°C. A partial fragment of the *B. dothidea Actin* gene was used to normalize the RNA samples for each qRT-PCR, and each treatment was conducted with three technical replicates. The absolute quantitative standard curves were established by qRT-PCR analysis of plasmids containing the corresponding BdcRNA cDNAs serially diluted to concentrations ranging from 10^5^ −10^10^ ng μL^−1^. The BdcRNA concentration (copies μL^−1^) was calculated based on the amount of cDNA as referred to a standard curve, which corresponded to equal amounts of BdcRNAs extracted from 1 mg mycelia.

### 4.10 Transcriptome and proteomic analysis

The fungal RNA extraction and transcriptome analysis were performed in triplicate by Beijing Novogene Bioinformatics Technology Co. Ltd. according to their pipeline. Briefly, sequencing libraries were generated using the NEBNext® UltraTM RNA Library Prep Kit for Illumina® (NEB, USA) with 1 μg of total RNAs, added index codes to attribute sequences to each sample, sequenced using an Illumina Novaseq platform and generated 150 bp paired-end reads. The raw paired-end reads were trimmed, quality controlled, and aligned to the reference genome using Hisat2 v2.0.5. DEG analysis was performed using the DESeq2 R package (1.16.1). Fungal protein extraction and proteomic analysis were performed by Jingjie PTM BioLab (Hangzhou) Co. Ltd. Briefly, fungal proteins were extracted, digested with trypsin, dissolved in 0.1% formic acid (solvent A), and directly loaded onto a home-made reverse-phase analytical column (15-cm length, 75 μm i.d.). The peptides were subjected to NSI source followed by tandem mass spectrometry (MS/MS) in Q ExactiveTM Plus (Thermo) coupled online to the UPLC.

GO enrichment analysis of DEG were implemented by the cluster Profiler R package, in which gene length bias was corrected. A *p*-value < 0.05 was considered to indicate statistically significant enrichment. The GO annotation proteome was derived from the UniProt-GOA database (http://www.ebi.ac.uk/GOA/). Cluster membership was visualized by a heat map using the “heatmap.2” function from the “gplots” R-package.

### 4.11 Statistical Analysis

Descriptive statistics were determined, and chi-square tests, one-way ANOVA and Tukey post-hoc tests were conducted using SPSS Statistics 17.0 (Win Wrap Basic; http://www.winwrap.com). A *p*-value ≤ 0.05 was considered to indicate significance. For the growth rate and virulence assays, the mean values for the biological replicates are presented as column charts with error bars representing SD. The graphs were produced in Excel (Microsoft) and GraphPad Prism 7 (GraphPad software).

## Acknowledgements

The authors thank the late Professor Ricardo Flores, Instituto de Biologia Molecular y Celular de Plantas, (IBMCP, UPV-CSIC), Valencia, Spain, for improving the manuscript. Profs. Daohong Jiang and Kenichi Tsuda, College of Plant Science and Technology, Huazhong Agricultural University, Wuhan, Hubei 430070, China, for kindly reviewing the manuscript, and Prof. Guotian Li, College of Plant Science and Technology, Huazhong Agricultural University, Wuhan, Hubei 430070, China, for kind contribution of the vector pBS-NEO-Histone-mCherry.

## Author Contributions

W.X. conceived this study, designed the investigation, wrote the manuscript, and supervised the project. K.D. conducted most of the experiments, C.X. assessed the BdcRNA1 transfection, monitored growth under different stress conditions and preformed co-infection with mycoviruses, R.L. conducted infection of other fungi, J.J cloned BdRV1, and L.K. participated in some data analysis. S.L., N.H. and G.W. participated in the design of the investigation. I.K.-L. and R.H.A.C. improved English, presentation, and discussion.

## Data and materials availability

Sequence data supporting the findings of this study has been deposited in GenBank under accession numbers MH371150-MH371155 for BdcRNAs, and KP245734-KP245738 for BdRV1, respectively. The remaining data are available within the article and the supplementary materials files and from the corresponding author upon request.

