## Supplementary figures for "Novel Viroid-like RNAs Naturally Infect a Filamentous Fungus"

Figure S1

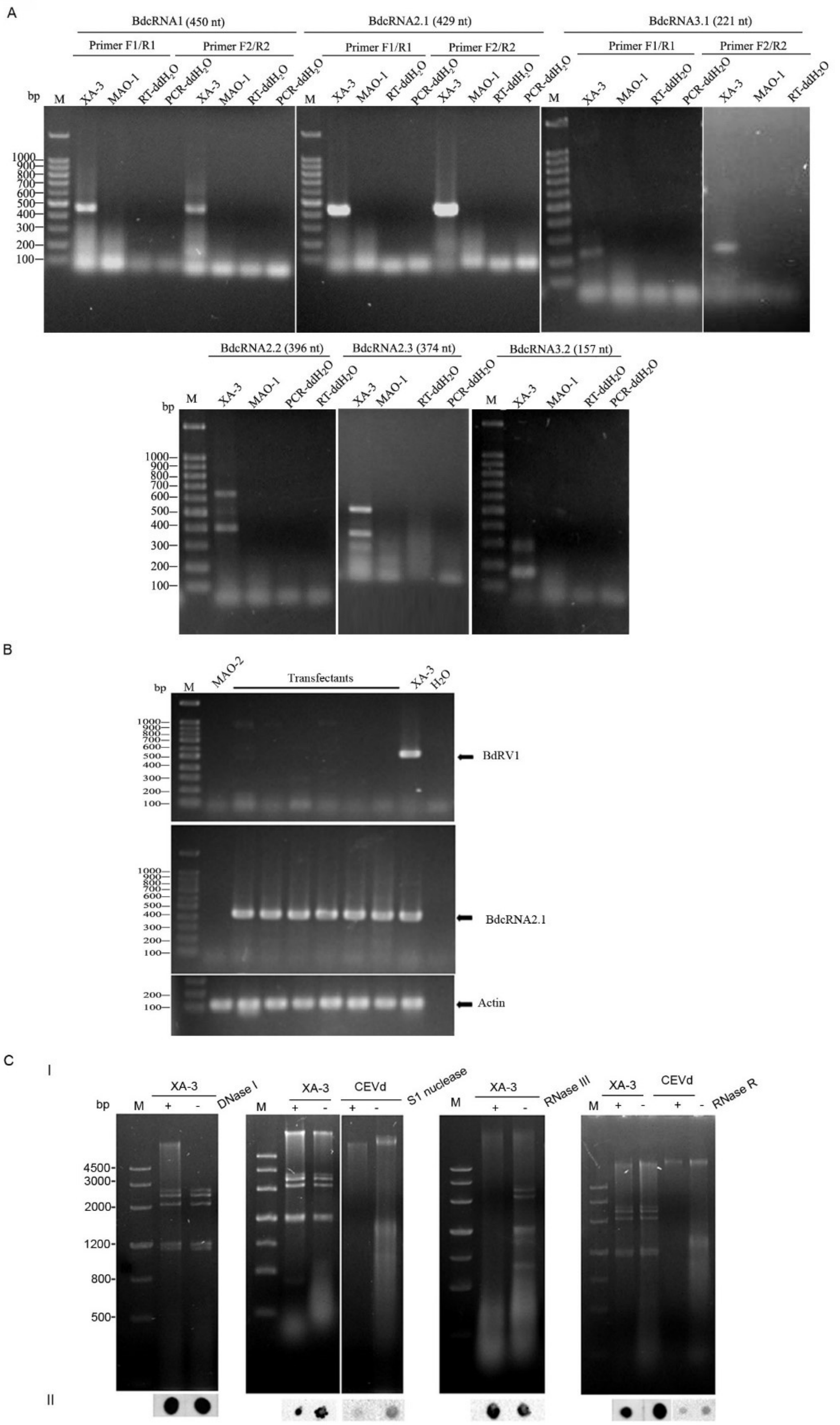


**Figure S1.** RT-PCR and dot blot detection of Botryosphaeria dothidea circular RNAs (BdcRNAs) and Botryosphaeria dothidea RNA virus 1 (BdRV1). (A) RT-PCR detection of BdcRNAs 1, 2.1, 2.2, 2.3, 3.1 and 3.2 using abutted primer pairs of opposite polarity (-F1 and -R1, Table S1) designed based on obtained contigs by assembling partial cDNAs amplified from each individually purified cRNAs or based on Sanger sequencing (-F2 and -R2). Target BdcRNAs are indicated above the gel. M, DNA size marker. (B) RT-PCR identification of BdRV1 together with BdcRNA2.1 and *Actin* gene in the protoplast-generating MAO-2 subisolates transfected with BdcRNAs. (C) Electrophoretic analysis on a 1.2% agarose gel of nucleic acid preparation from XA-3 before (−) and after (+) digestion with DNase I, S1 nuclease, RNase III and RNase R (I); dot-blot of the nucleic acid preparation with the BdcRNA2 antisense probe (II). Nucleic acid preparation of citrus exocortis viroid (CEVd), a circular ssRNA, was included in parallel as control and detected using the CEVd.188 antisense probe.

Figure S2

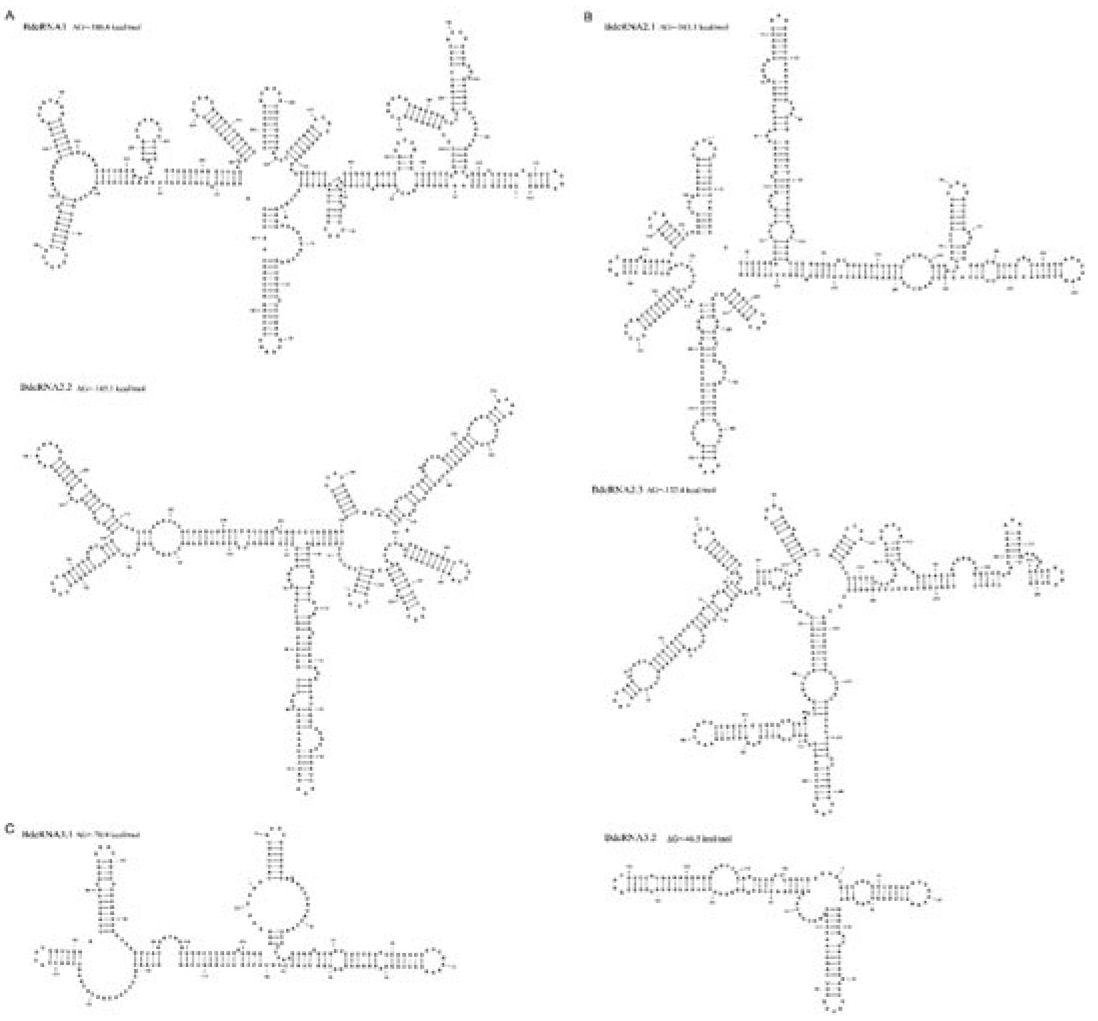


**Figure S2.** Predicted secondary structures of BdcRNAs in the lowest energy determined using the RNA Structure prediction tool in CLC RNA Workbench software (Version 4.8, CLC bio A/S). (A-C) The secondary structures of BdcRNAs 1 (A), 2.1, 2.2, 2.3 (B), 3.1 and 3.2 (C), respectively.

Figure S3

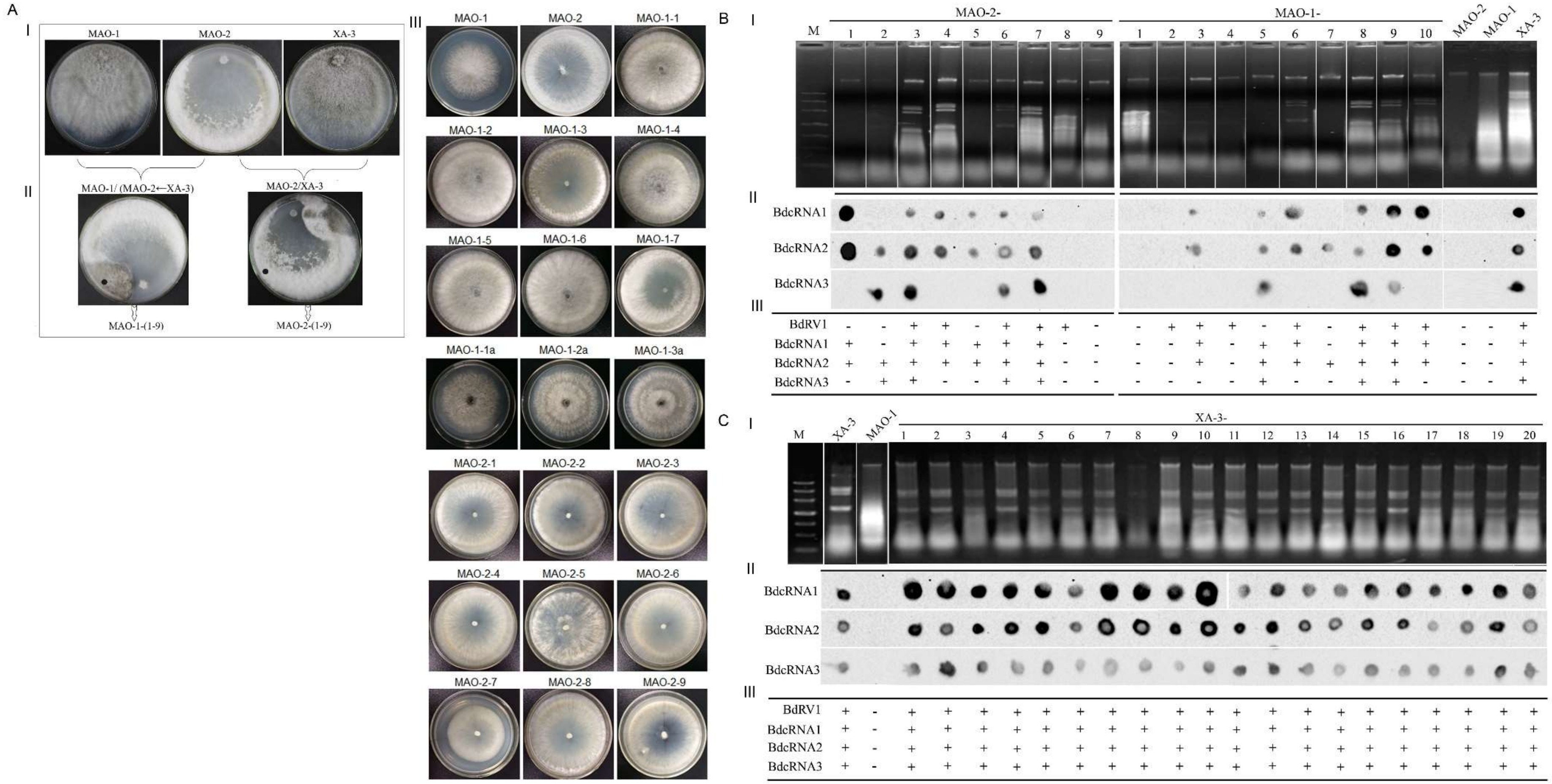
A
I
MAO-1
MAO-2
XA-3
II
MAO-1/ (MAO-2←XA-3)
MAO-2/XA-3
MAO-1-(1-9)
MAO-2-(1-9)
III
MAO-1
MAO-2
MAO-1-1
MAO-1-2
MAO-1-3
MAO-1-4
MAO-1-5
MAO-1-6
MAO-1-7
MAO-1-1a
MAO-1-2a
MAO-1-3a
MAO-2-1
MAO-2-2
MAO-2-3
MAO-2-4
MAO-2-5
MAO-2-6
MAO-2-7
MAO-2-8
MAO-2-9
B
I
M
MAO-2-
MAO-1-
MAO-2
MAO-1
XA-3
II
BdcRNA1
BdcRNA2
BdcRNA3
III
BdRV1
BdcRNA1
BdcRNA2
BdcRNA3
C
I
M
XA-3
MAO-1
XA-3-
II
BdcRNA1
BdcRNA2
BdcRNA3
III
BdRV1
BdcRNA1
BdcRNA2
BdcRNA3

**Figure S3** Horizontal transmission analysis of BdcRNAs and colony appearance of the subisolates. (a) Colony appearance of the strains involved in the contact cultures. Colonies of strains MAO-1, MAO-2 and XA-3 in single culture (I), contact culture (II), and resulted subisolates (III). The "●" indicates the location where a mycelial agar plug was removed for to generate a subisolate derived from strain MAO-1 or MAO-2. (b) and (c) Nucleic acid preparations (I), dot-blot analysis of BdcRNAs 1 to 3 (II), summary table of the infection of BdcRNAs and BdRV1 (III) of subisolates derived from the contact cultures (b) and strain XA-3 conidia (c), respectively. The "+" and "-" indicate the presence and absence of BdRV1 or BdcRNAs, based on dsRNA detection by 1.2% agarose gel electrophoresis (BdRV1) and dot blots (BdcRNAs), respectively. M, DNA size marker. (d) Colony appearance of the subisolates of MAO-2 transfected with mixed BdcRNAs eluted from gel delimited by the DNA markers of 100 and 500 bp.

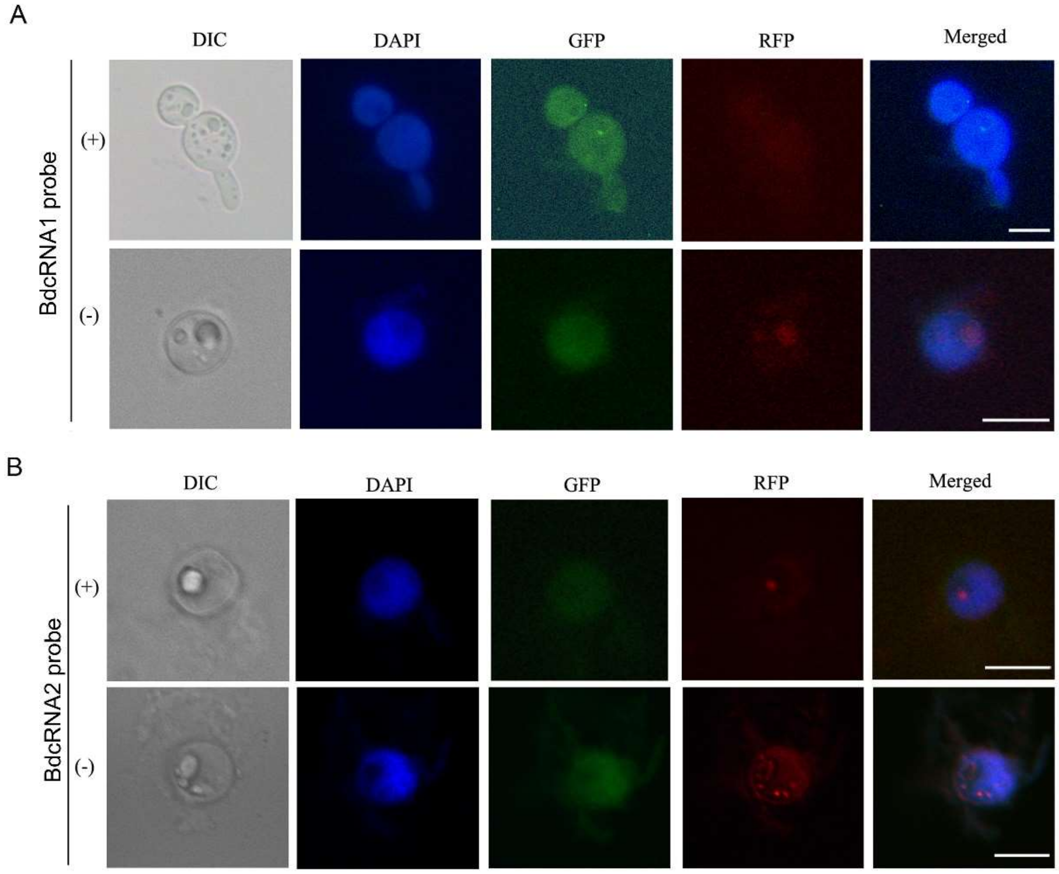


**Figure S4.** Subcellular location analysis of BdcRNAs. (A and B) FISH to detect subcellular location of plus (+) and minus (-) strands of BdcRNA1 (A) and BdcRNA2.1 (B) in protoplasts of strain XA-3 by Alexa Fluor 488–labeled riboprobes (green fluorescence).The nuclear was indicated with mCherry, binding the fungal nucleosome (red fluorescence), and stained with DAPI in blue color in higher concentration than in cytoplasm.

Figure S5

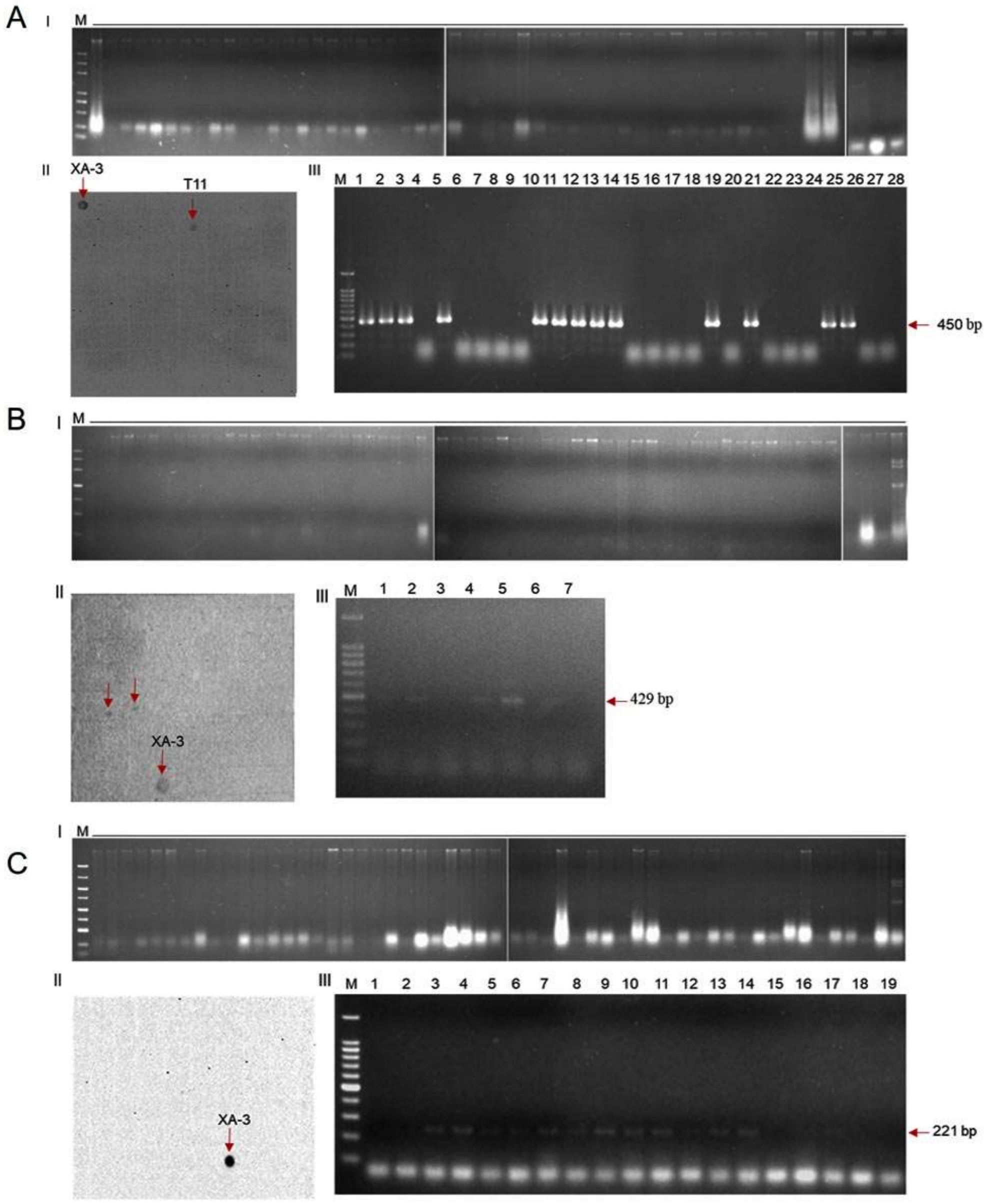


**Figure S5**. Detection of BdcRNAs in protoplast-generated colonies derived from *Botryosphaeria dothidea* MAO-2 transfected with dimeric RNAs that were transcribed from BdCRNA dimeric cDNAs inserted in pGEM-T after digested with *Nde* I. Total 50, 56 and 52 colonies were detected for the transfectants derived from fungal strain transfected with BdCRNAs 1 (A), 2.1 (B) and 3.1 (C), respectively, using dot blot (II) and RT-PCR (III) based on the extracted nucleic acids using column methods (I).

Figure S6

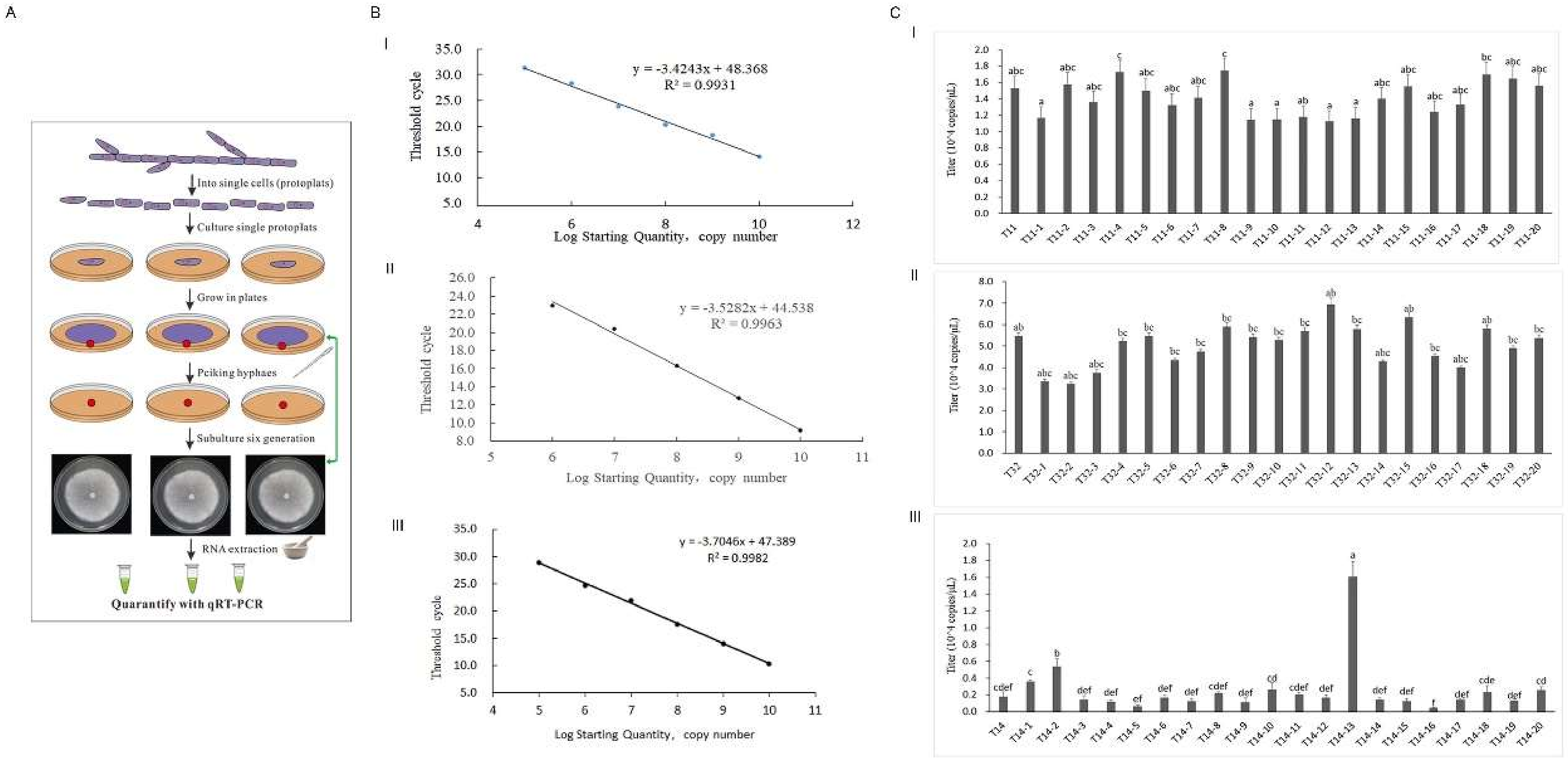
A
Into single cells (protoplats)
Culture single protoplats
Grow in plates
Peiking hyphaes
Subulture six generation
RNA extraction
Quarantify with qRT-PCR
B
I
y = -3.4243x + 48.368
R² = 0.9931
Threshold cycle
Log Starting Quantity, copy number
II
y = -3.5282x + 44.538
R² = 0.9963
Threshold cycle
Log Starting Quantity, copy number
III
y = -3.7046x + 47.389
R² = 0.9982
Threshold cycle
Log Starting Quantity, copy number
C
I
Titer (10^4 copies/μL)
II
Titer (10^4 copies/μL)
III
Titer (10^4 copies/μL)

**Figure S6** Quantitative analysis of the systematic infection of BdcRNAs in transfectants. (A) Flow chart for the qRT-PCR analysis of the systematic infection and stability of BdcRNAs in individual cells after sub-culture for six generations. (B) The standard curve for qRT-PCR analysis of cDNA plasmids of BdcRNA1 (I), 2.1 (II) and 3.1 (III) after serially diluted ranging from $10^5$ to$10^{10}$ ng/μL. (C) Bar graph for the BdcRNA titers for twenty individual protoplast cells after sub-cultured for six generation for the transfectants of MAO-2 transfected by BdcRNA1 (T11; I), BdcRNA2.1 (T32; II), and BdcRNA3.1 (T14; III). The cell-generated subisolates are serially termed -1 to -20.

Figure S7

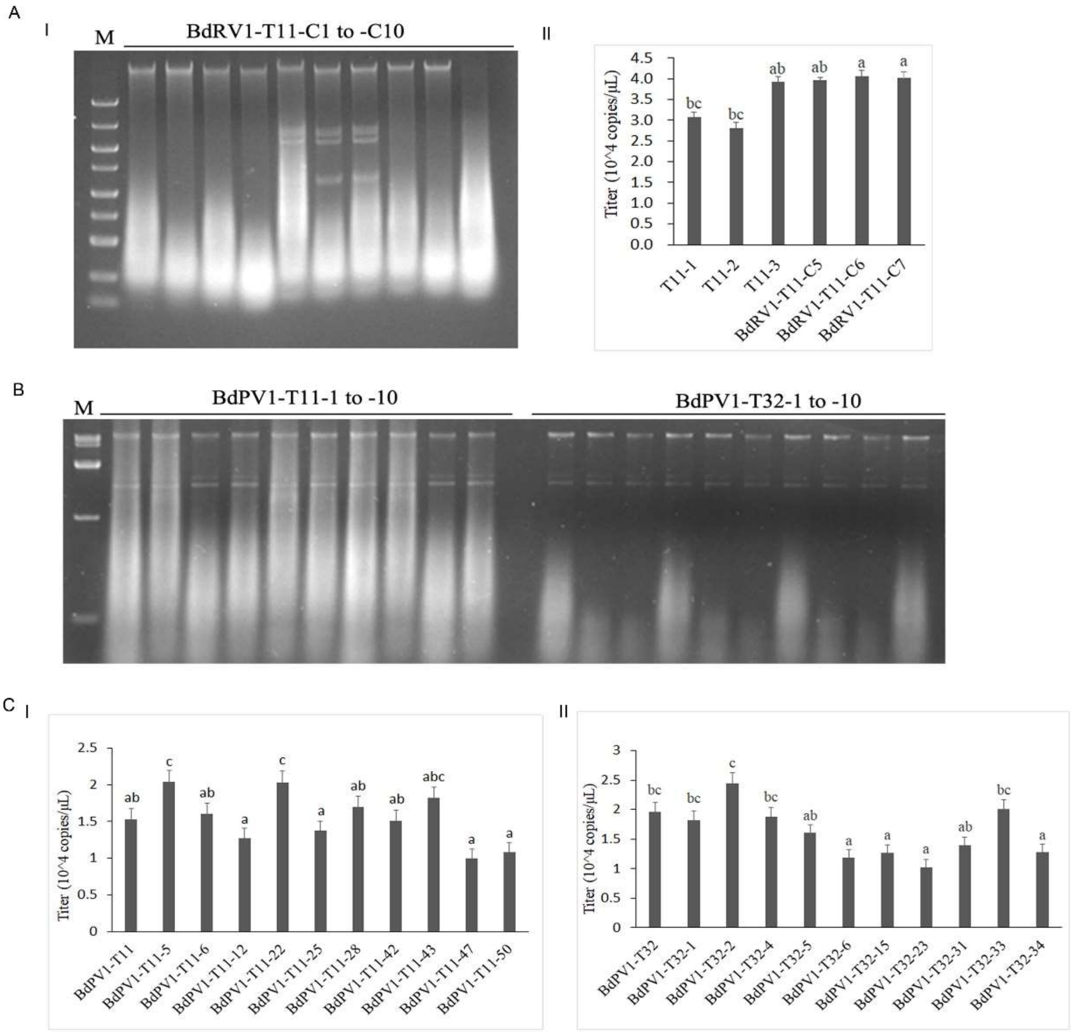


**Figure S7** Quantitative analysis of BdcRNA replication affected by mycovirus. (A) DsRNA detection of ten subisolates of T11 infected by Botryosphaeria dothidea RNA virus 1 (BdRV1) after contact culture with MAO-2-9 (containing only BdRV1) (I), and qRT-PCR analysis of BdcRNA1 titers in the subisolates before (termed T11-1 to -3) and after (BdRV1-T11-C5 to C7) co-infected with BdRV1 (II). (B) DsRNA detection of ten subisolates of T11 and T32 after transfected by Botryosphaeria dothidea partitivirus 1 (BdPV1) virions. (C) qRT-PCR analysis of BdcRNA1 (I) and BdcRNA2.1 (II) titers in the subisolates of T11 and T32, respectively, after infected by BdPV1 and sub-culture for six generation. M, DNA size marker.

Figure S8

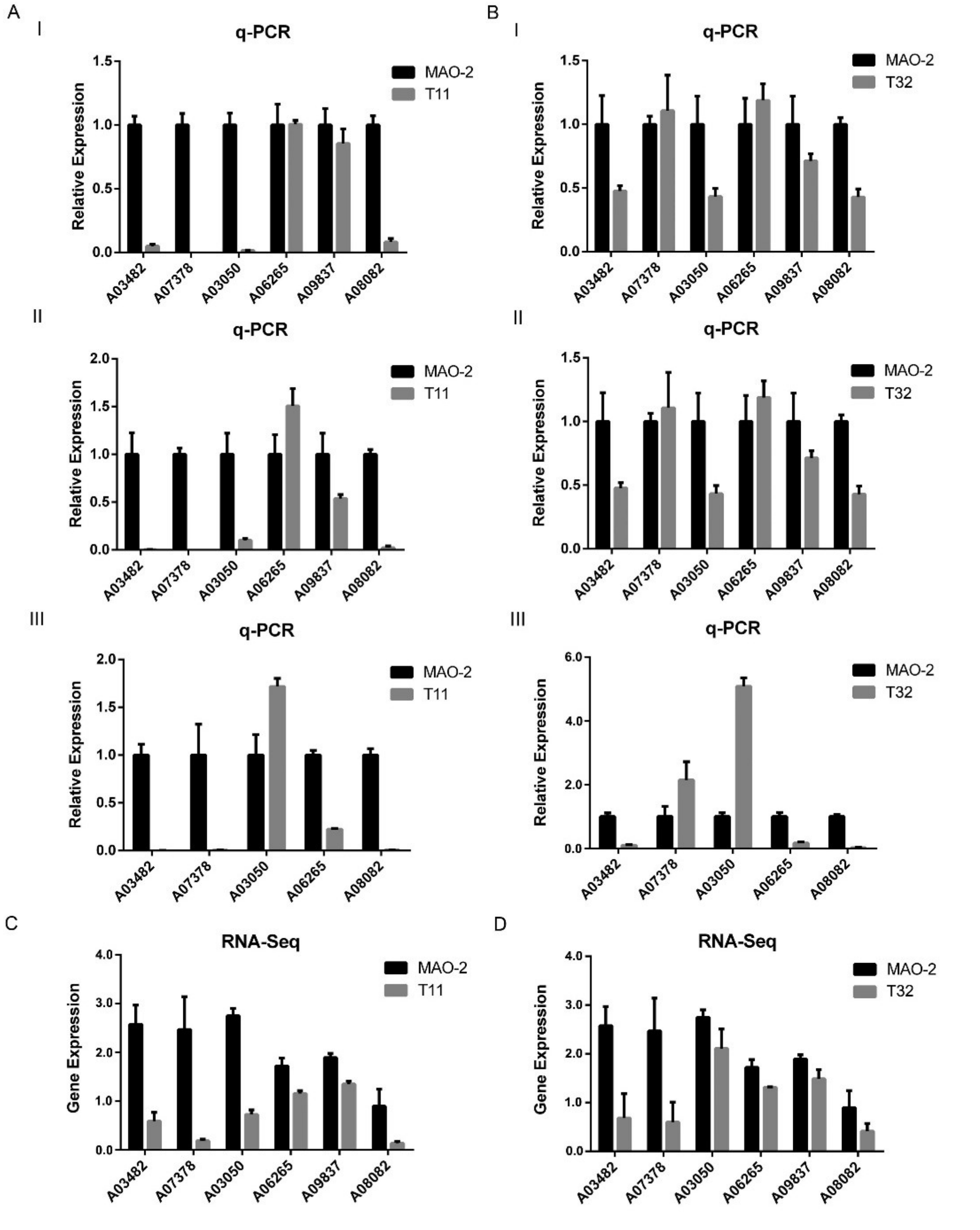


**Figure S8** Quantitative real-time PCR (qRT-PCR) analysis of the relative expression changes of six genes in MAO-2 and BdcRNA1 and 2.1 transfectants (T11 and T32, respectively). QRT-PCR analyses were conducted after cultured at 6 (I), 8 (II) and 10 (III) dpi on PDA, for T11 (a) and T32 transfectants (b) as compared with MAO-2, respectively. The transcriptome sequencing data of mycelia collected at 8 dpi on PDA were involved for comparison for T11 (C) and T32 (D), respectively.
